# *cryoWEight*: Conformational ensembles from cryo-EM particle images with weighted ensemble simulations

**DOI:** 10.64898/2026.09.28.755042

**Authors:** Anupam Anand Ojha, Praveen Ranganath Prabhakar, Ioan Andricioaei, Sonya M. Hanson

## Abstract

Atomistic simulations often struggle to capture the full conformational landscape of biomolecular complexes on experimentally relevant timescales, leaving gaps in our understanding of functionally relevant thermodynamics and kinetics. Cryo-electron microscopy (cryo-EM) provides high-resolution structural snapshots spanning multiple conformational states, but translating these static images into dynamic mechanisms remains challenging. Existing approaches for integrating cryo-EM data with molecular simulations either perturb the underlying dynamics through artificial biasing potentials or remain confined to conformations already present in the prior ensemble. We introduce *cryoWEight*, an automated and extensible Python framework that couples iterative weighted ensemble (WE) simulations with ensemble reweighting against cryo-EM particle images to steer simulation trajectories towards conformational states consistent with cryo-EM populations. We have found that reweighting in the image space introduces a directional bias towards the target distribution even when the prior ensemble has no overlap with the target distribution in the collective variable space. The reweighting requires only the state likelihoods that set the ensemble weights, and any observable that defines these likelihoods can serve as the target distribution, therefore generalizing beyond the application of cryo-EM. *cryoWEight* is validated on three proteins: chignolin, the N-terminal domain of ribosomal protein L9, and adenylate kinase. The framework converges towards the target conformational states across different sampling strategies and noise levels, providing a scalable route to integrate cryo-EM data with enhanced sampling approaches.

## I. Introduction

Cryo-electron microscopy (cryo-EM) has emerged as a leading technique in structural biology to resolve the three-dimensional (3D) structures of biomolecules at high resolution^1–5^. With technological advancements in instrumentation^6,7^ and computationally efficient algorithms^8,9^, cryo-EM has successfully addressed challenging questions related to biophysical phenomena such as protein folding^10^, ligand binding^11,12^, and functionally relevant conformational transitions^13,14^ by providing high-resolution structural snapshots of biomolecules across multiple conformational states^15^. However, translating these static snapshots into dynamic mechanisms, thermodynamic properties, and kinetic rates remains challenging and will require integration with complementary approaches, such as all-atom molecular dynamics (MD) simulation^16–18^. While MD simulations have been routinely implemented to probe biomolecular behavior at microscopic spatio-temporal resolution^19–21^, high free energy barriers and kinetic traps leave the equilibrium populations of conformational states difficult to resolve by unbiased MD^22,23^, thereby rendering it advantageous to be coupled with an external source of structural information to guide simulations toward inaccessible states. Thus, the use of the molecular snapshots across conformational states available in cryo-EM data to enhance efficient MD sampling methods is a natural complement.

MD simulations of rare events, including large-scale protein conformational transitions^21,24,25^, folding/unfolding^26,27^, and ligand binding^28–30^, remain computationally expensive because biomolecules reside in metastable basins for extended periods. Efficient sampling of such transitions therefore requires methods that redistribute computational effort towards rarely visited regions of the configuration space^31–34^. The weighted ensemble (WE) method^35,36^ stands out as an efficient approach for exploring conformational space, in which parallel trajectories are periodically split and merged without introducing external biases^37^. Since its foundation^35^, the WE method has been successfully implemented on challenging rare event problems such as protein (un)folding^38–41^, protein-protein binding^42^, membrane permeation and peptide translocation^43,44^, ligand binding to nucleic acids^45^, cryptic pocket opening^46^, viral glycoprotein opening^47^, and proteinligand unbinding^48^. Significant advancements in the implementation of the method have also been reported in a recent review^49^. However, extended simulation runs often suffer from weight degeneracy, in which successive resampling cycles concentrate probability on a few walkers and collapse the effective sample size, thereby limiting exploration of new conformational space^48,50^. Several reweighting strategies have been proposed to counter this, including equilibrium and steady-state reweighting^51,52^, history-augmented Markov state models^53,54^, and trajectory stratification^55^, alongside transition-based reweighting analysis developed for data collected across multiple thermodynamic states^56,57^. All of these strategies, however, reweight against the simulation history itself. Because unvisited conformations carry zero weight, no reweighting operation can assign them a non-zero probability, and the corrected ensemble can never extend beyond the states already sampled. Overcoming this limitation can be possible with a reference that constrains the equilibrium populations of states the simulation has not itself reached.

While this reference can be provided from a variety of sources, cryo-EM provides an experimental conformational distribution that is ideally suited to this task, and its integration with molecular simulations offers a natural path forward. Methods for extracting conformational heterogeneity from cryo-EM images are still in active development. Most methods infer this heterogeneity from cryo-EM images alone, without coupling to simulation^58^. These methods range from classifying particles into discrete states via maximum-likelihood methods^59,60^ to representing continuous latent spaces via machine learning approaches^61^. One of the first methods to extract continuous landscapes was the manifold embedding (ManifoldEM) approach to heterogeneity in cryo-EM^62–66^. Another approach uses linear subspace methods, which represent conformational heterogeneity in a low-dimensional space of eigenvolumes, obtained either by eigenanalysis of the covariance of the 3D volumes^67^ or by fitting the subspace directly from the images without forming that covariance^68,69^, with RECOVAR combining a regularized covariance estimator with adaptive kernel regression^70^. Machine learning approaches, including cryoDRGN^61^, 3DFlex^71^, DynaMight^72^, and cryoSTAR^73^, model continuous conformational heterogeneity directly from the images. More recently, approaches that combine molecular simulation to access the heterogeneity in cryo-EM data have been developed. Most notably for this work, ensemble reweighting methods couple cryo-EM images to molecular simulation, assigning weights to simulated conformations using Bayesian likelihoods computed on individual particle images^74–78^. There are also metainference approaches which take a similar approach but use Bayesian likelihoods against reconstructed density maps^79–82^. A recent method fits atomic coordinates to individual particle images under a steering restraint^83^, returning static structures whose populations are determined by the restraint rather than by Boltzmann statistics. Benchmark datasets with known ground truth and direct comparisons against MD simulations now quantify the accuracy of these approaches^84–86^. The recovered populations either remain confined to conformations already present in the input ensemble or the observed particles, or are obtained under restraints that bias the underlying dynamics away from the Boltzmann distribution.

Despite such methodological advances, existing approaches for integrating cryo-EM data with molecular simulations either perturb the underlying dynamics through artificial biasing potentials or remain constrained to conformations already present in the simulated ensemble. The ability to simultaneously generate new conformations in undersampled regions and correct population biases against cryo-EM distributions, therefore, remains an unmet need. We present *cryoWEight*, an automated and extensible Python framework that couples iterative enhanced sampling with cryo-EM ensemble reweighting in a closed feedback loop, steering simulation trajectories towards equilibrium populations consistent with the particle images (Figure 1a). The generalized protocol clusters an initial ensemble from molecular dynamics, coarse-grained simulation, or structure prediction models^87–89^ in the collective variable (CV) space, reweights the representative structures against the particle images, and seeds the next round of conformational space exploration under enhanced sampling, iterating until the populations converge to the cryo-EM distribution (Figure 1b). The current framework is implemented with a WE sampling scheme. The cryo-EM signal periodically adjusts walker weights by their likelihood of generating the observed particle images and corrects population biases without altering the underlying dynamics. The reweighted conformations then seed the subsequent WE simulation and direct the sampling towards previously inaccessible regions. This feedback loop drives the ensemble to progressively converge towards the cryo-EM distribution by generating new conformations guided by the experimental signal. Ensemble reweighting depends on the target distribution only through the likelihood assigned to each conformation and not on the observable from which that likelihood is computed. The principle underlying *cryoWEight* therefore holds for targets defined by observables beyond cryo-EM particle images, and extends to any distribution expressible as a likelihood over structures^90,91^. The *cryoWEight* framework is validated on three proteins of increasing complexity, namely the 10-residue miniprotein chignolin in explicit solvent, the 39-residue N-terminal domain of ribosomal protein L9 (NTL9) in implicit solvent, and adenylate kinase (ADK) in explicit solvent, which undergoes large-scale rearrangements of its nucleoside monophosphate (NMP) and LID domains between open and closed catalytic states.

**FIG. 1:**
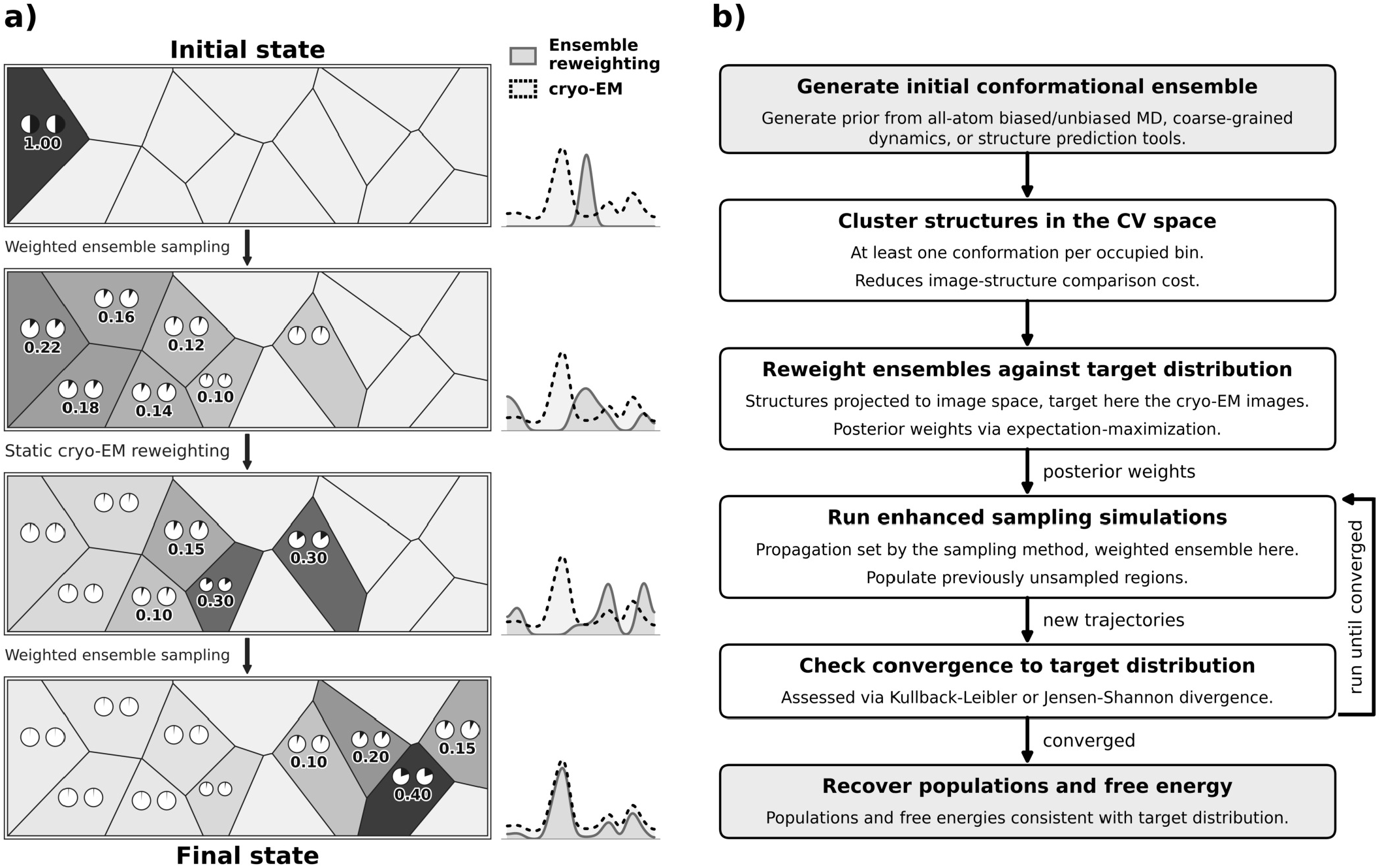
(a) Iterative workflow of weighted ensemble (WE) sampling and cryo-EM reweighting on a 2D free energy landscape. Each walker is drawn as a circle within a Voronoi cell with a shaded sector proportional to its statistical weight, and each cell is shaded by its cumulative weight. The 1D profiles beside each panel show the per-bin distributions, with the reweighted ensemble shown as the light gray curve and the cryo-EM target as the dark gray curve. WE sampling propagates walkers by splitting and merging. Cryo-EM reweighting then redistributes their weights according to the likelihood of reproducing the observed particle images, and the reweighted ensemble seeds the next iteration. Over successive cycles, the distribution converges onto the cryo-EM distribution. (b) Generalized *cryoWEight* protocol. An initial ensemble from all-atom molecular dynamics, coarse-grained simulations, or a structure prediction model is clustered in the collective variable space into representative structures, which are then reweighted against the target distribution. The posterior weights seed the next set of simulations to populate unsampled regions. The procedure terminates once equilibrium populations and free energies consistent with the target distribution are recovered.

## II. Theory and Methods

This section presents the methodological framework of *cryoWEight*. Section II A introduces the weighted ensemble method for conformational sampling, Section II B describes the ensemble reweighting methods that recover equilibrium populations from cryo-EM particle images, Section II C details the *cryoWEight* framework that integrates WE sampling with cryo-EM reweighting, and Section II D provides the simulation setup for chignolin, NTL9, and ADK.

### A.Weighted ensemble simulations

The WE method^35^ runs an ensemble of parallel trajectories, each with a statistical weight that contributes to the overall probability distribution. During simulation, trajectories in the undersampled regions of the configuration space are replicated (split), while those in the well-sampled regions are stochastically pruned (merged). The splitting and merging operations conserve the total probability, and the underlying dynamics remain unbiased as no external forces modify the free energy landscape, ensuring that the simulations generate statistically exact estimates of both equilibrium and kinetic properties^37^. The mathematical framework governing WE simulations is described below.

#### 1. Initialization

The CV space is first discretized into a set of non-overlapping bins (e.g., rectilinear cells). WE simulations are typically initiated by placing replicas of a single molecular configuration into one bin corresponding to a metastable state, with each replica assigned a statistical weight. Alternatively, multiple conformational states can be employed^92^, with a fixed set of replicas placed in each bin. The corresponding weights are then assigned according to a chosen or arbitrary initialization scheme. For estimating kinetic properties, WE simulations are carried out with recycling boundary conditions, in which the source and sink states are predefined. At the steady state, the flux per unit time from the source to the sink is computed to estimate mean first passage times via the Hill relation^36,51,52,93^. For equilibrium sampling, simulations proceed without recycling, and properties such as free energies along the progress coordinate are estimated from the relative weights of configurations in the ensemble.

Let initial conformations construct an ensemble of *M*_w_ trajectories (or walkers), each with an assigned statistical weight. This ensemble represents an initial probability distribution over the configuration space Ω. Let us consider *K* initial configurations 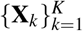, each associated with a probability *p*_*k*_ such that 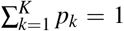. For each configuration *k, M*_*k*_ walkers are initialized such that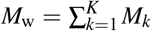 and each walker *i* started from *k* is assigned weight 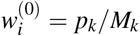 The initial probability distribution over configurations is

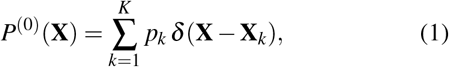

where *δ* denotes the Dirac delta function. The distribution *P*^(0)^ is normalized since

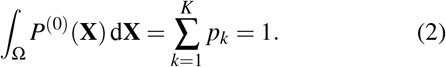

#### 2. Collective variable mapping and bin assignment

Each configuration X ∈ Ω is mapped to a *d*-dimensional CV vector **q = q(X)** ∈ ℝ^*d*^. The CV space is partitioned into *N* non-overlapping bins 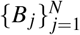with 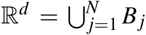 and 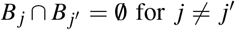. Each bin is a rectilinear cell of the CV space. For walker *i* at the zeroth iteration, the CV is 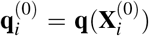 and the walker is assigned to the bin that contains 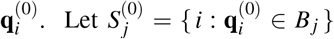 denote the set of walkers in bin *B*_*j*_ at iteration 0. The total weight in bin *j* is 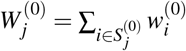 and the number of walkers is 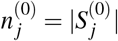 The bin weights satisfy

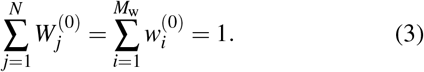

Given that the conformational space is represented by two-dimensional (2D) CVs, the mapping is expressed as q = (*q*^(1)^, *q*^(2)^), and each dimension is discretized into *n*_1_ and *n*_2_ bins, respectively. The bin 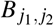 is

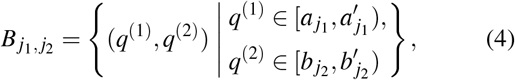

where 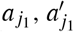 and 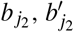 are the boundaries in the first and second coordinate for *j*_1_ = 1, …, *n*_1_ and *j*_2_ = 1, …, *n*_2_, and the *N* bins of the discretized CV space correspond to *N* = *n*_1_*n*_2_. The bins are adaptively refined by the minimal adaptive binning (MAB) mapper^94^, and the resulting boundaries are recomputed at each iteration. Along each progress coordinate, the MAB mapper distributes a fixed number of evenly spaced bins between the trailing and leading walkers, each of which is also assigned a bin. Additional bins are then allocated to the walkers maximizing 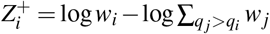and 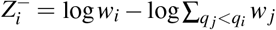 where *q*_*i*_ is the progress coordinate of walker *i*^94^. The mapper is nested within a single outer bin of the rectilinear partition, while the remaining walkers are binned solely by the partition. The location of the outer bin is a parameter of the binning scheme and remains constant as the ensemble advances along the progress coordinate, being updated only between successive simulations.

#### 3. Propagation and resampling

The WE simulation proceeds iteratively through cycles of propagation and resampling (splitting and merging). At each iteration *n*, each walker *i* is propagated independently for a fixed resampling time *τ* via unbiased stochastic dynamics (e.g., Langevin or Brownian dynamics), leading to a new configuration 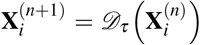 Here, 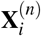 denotes the configuration at the start of iteration *n* and *D*_*τ*_ is the stochastic propagator for the interval *τ*. The new configuration is then mapped to the CV space as 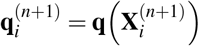,which determines the bin assignment 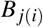 with 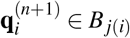.After propagation, resampling ensures that the number of walkers within each bin matches a pre-defined target (e.g., *M*_bin_ walkers per bin) while conserving the total statistical weight. Let 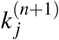 be the number of walkers in bin *B*_*j*_ after propagation, and *M*_bin_ be the target number of walkers per bin. If 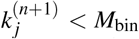 walkers in *B*_*j*_ are split until the count reaches *M*_bin_. When a walker *i* is split into *m*_*i*_ copies, each child receives weight 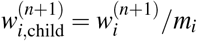 such that

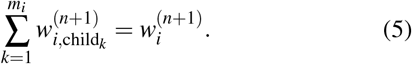

Conversely, 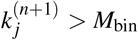 walkers are pruned or merged, and the pruned weights are stochastically reassigned to survivors such that

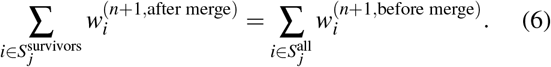

After resampling, the ensemble remains normalized such that 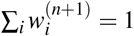 The weight for each bin after resampling is

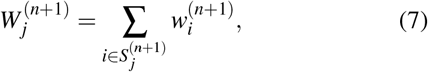

where 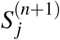 is the set of walkers in bin *B*_*j*_ after resampling.

### B. Cryo-EM ensemble reweighting

While WE simulations provide enhanced sampling of conformational space^36^, the resulting ensemble may still deviate from the true equilibrium distribution due to inaccuracies in the force field^95–97^, incomplete sampling convergence^98–101^, or biases introduced by the choice of CVs and resampling strategy^38,48,102^. Cryo-EM particle images are single-molecule snapshots of individual conformations rather than ensemble averages. Conformational distributions are therefore extracted directly from the images by Bayesian inference^74,76,77^. Building on the ensemble reweighting formalism, the conformational weights are inferred by an expectation-maximization algorithm^8,59,103^. The reweighting is performed in image space rather than in the CV space, ensuring that even conformations with poor CV overlap can be discriminated by their distinct 2D projections^76^. The reweighting procedure is described below.

#### 1. Clustering and selection of representative structures

Let us consider an MD trajectory with *T* frames 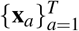, each given a uniform initial weight 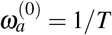 with 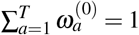. For each frame *a*, CVs are computed as

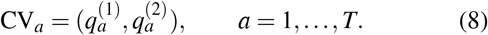

The conformational space is partitioned into a regular 2D grid of bin width Δ with

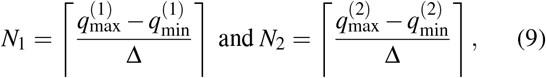

and bin edges

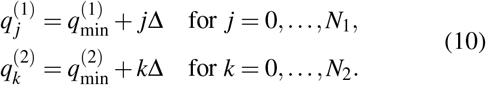

Each frame *a* is assigned to a bin (*u, v*) determined by

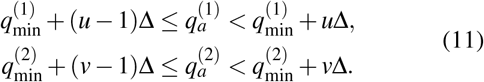

Let *S*_*u,v*_ denote the set of indices of the frames assigned to bin (*u, v*) through equation 11. For each occupied bin (*u, v*), a representative frame *f*_*u,v*_ is selected from the members of *S*_*u,v*_, which differ by less than the bin width Δ along each CV. The weights of the frames within the bin are merged into this representative, which is then assigned the total weight of the bin,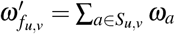 and the remaining frames are set to 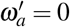for *a* ∉ {*f*_*u,v*_}. The merged weights are then normalized as 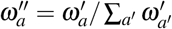 with 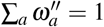 set of *M* non-redundant representative frames 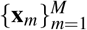 (with *M* equal to the number of occupied bins) is thus obtained, each assigned a normalized weight 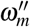. In the prior ensemble reweighting work^77^, the conformations are clustered by k-medoids on the pair-wise C*α*-RMSD matrix, with the cluster medoids as the representative structures. The grid partitioning in *cryoWEight* instead selects representative structures within the user-defined CV space, thereby reducing the number of structures entering the image likelihood. The WE bins and the clustering grid are distinct partitions of the same conformational space. The WE bins are defined by the rectilinear partition with the nested MAB mapper (Section II A) and specify the regions of the conformational space within which walkers are split and merged to the target count during resampling. The clustering grid merges the conformations within each occupied bin into a single representative conformation at the bin width resolution. The representative frames now serve as the input ensemble for reweighting against cryo-EM images.

#### 2. Ensemble reweighting

The reweighting step in *cryoWEight* adapts the algorithm of the prior ensemble reweighting work^77^, where the forward model for image generation, the calculation of the distance between simulated and observed images, and the estimation of the global noise level are retained, while the Markov chain Monte Carlo sampling of the weight posterior is replaced with an expectation-maximization optimization of the image likelihood. The current optimization step adjusts the statistical weights of the representative structures such that their projected images reproduce the observed particle images, thereby recovering the populations more reliably than counting-based estimates, which are sensitive to image noise^104^. Synthetic cryo-EM images are generated from a reference MD ensemble, and a template image *I*_*m*_ is simulated for each representative conformation x_*m*_ at the known pose of each particle image for the likelihood evaluation. In the forward model, each residue of a conformation is projected onto the image plane as a 2D Gaussian centered on its C*α* atom,

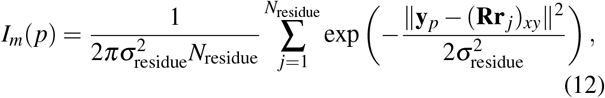

where *σ*_residue_ is the width of the Gaussian placed on each residue, **r**_*j*_ is the position of the C*α* atom of residue *j, N*_residue_ is the number of residues, **R** is the rotation matrix that defines the projection direction, y_*p*_ is the center of pixel *p*, and (**Rr**_*j*_)_*xy*_ denotes the in-plane components of the rotated coordinate. The contrast transfer function (CTF) is incorporated into the projected images in Fourier space, and pixelwise Gaussian noise of standard deviation 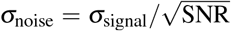 is added to the synthetic cryo-EM images, where *σ*_signal_ is the root mean square intensity of the noise-free image within a centered circular mask and the signal-to-noise ratio (SNR) determines the noise strength (Supplementary Tables S1-S3).

Given *M* representative conformations 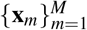 with initial weights 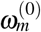 and *N* cryo-EM images 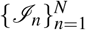, the pixelwise squared distance between the template of x_*m*_ and the cryo-EM image *I*_*n*_ is

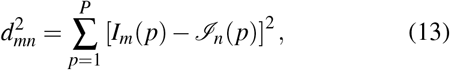

where *P* is the number of pixels. The template *I*_*m*_ in equation 13 is generated at the known pose and with the CTF of the particle image *I*_*n*_, and its mean intensity is subtracted before the distance is evaluated. Under a Gaussian noise model with the noise standard deviation 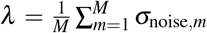averaged over one noise-free synthetic image per representative conformation, the log-likelihood for each structure-image pair is

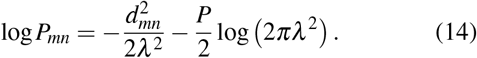

The normalization term in equation 14 is independent of the structure index *m* and cancels in the expectation step. The weights are optimized on the probability simplex by the expectation-maximization algorithm^77,104^. At iteration *t*, the expectation step computes the posterior probability 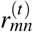 that image *n* was generated by structure *m*,

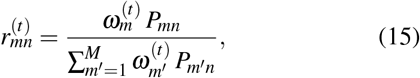

and the maximization step updates each weight as the image-averaged posterior probability,

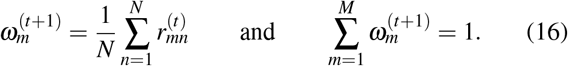

The iteration proceeds until the weights and the average log-likelihood

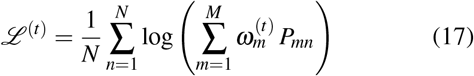

stabilize. Since clustering collapses each occupied bin (*u, v*) to a representative frame *f*_*u,v*_ with member set *S*_*u,v*_, the optimized weight 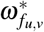 is redistributed uniformly to the original frames, 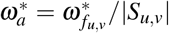 for *a* ∈ *S*_*u,v*_, remaining normalized so that 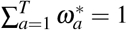.

The expectation-maximization algorithm has a unique fixed point on the probability simplex of non-negative weights that sum to unity. Upon convergence, the weights satisfy the self-consistency condition 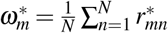 for *m* = 1, …, *M*, such that each optimal weight equals the average posterior probability across all images. The expectation-maximization algorithm guarantees that the average log-likelihood (equation 17) increases monotonically until a stationary point is reached, with the posterior probabilities *r*_*mn*_ serving as sufficient statistics for the reweighting update. Since the Gaussian likelihood (equation 14) is strictly positive for any finite *λ*, every conformation assigns a non-zero probability to every image, and no structure is ever excluded *a priori* from contributing to the reweighting. A smaller *λ* increases sensitivity to differences in image space distances, while a larger *λ* reduces it.

### C. *cryoWEight*: Iterative cryo-EM ensemble reweighting

The preceding sections describe WE simulations and cryo-EM ensemble reweighting as independent procedures. However, implementing them in isolation has limitations. Standalone reweighting can only redistribute probability among conformations already present in the ensemble and cannot generate new structures in undersampled regions of the configurational space. On the other hand, extended WE simulations may suffer from weight degeneracy, in which successive resampling cycles cause walker weights to diminish, thereby limiting exploration of new conformational states. To address these limitations, an iterative protocol is proposed that integrates WE simulations with cryo-EM reweighting in a feedback loop (Figures 1a and 1b, Supplementary Figure S1), where a forward model maps simulation trajectories to cryo-EM images for comparison with the target distribution. At each iteration, the reweighted ensemble seeds the subsequent WE simulation, biasing the sampling towards the cryo-EM distribution. The new set of conformations generated by the subsequent sampling is then reweighted against the cryo-EM images, and the cycle repeats until convergence. This strategy ensures that the ensemble progressively aligns with the experimental distribution while maintaining the continuous exploration of the conformational landscape. Throughout, an iteration refers to one cycle of the *cryoWEight* protocol, and a subiteration to one WE cycle of propagation and resampling within it.

#### 1. Clustering and selection of representative structures

Let ℓ denote the iteration index of the *cryoWEight* protocol.

The input at iteration ℓ is the ensemble 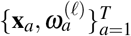, obtained from an unbiased MD simulation at ℓ = 0 or from the preceding WE simulation at ℓ > 0. The walker weights of the preceding simulation are not carried forward, and each frame is assigned an equal weight 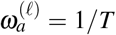 at the start of every iteration, preventing the accumulated weight degeneracy from propagating between iterations. CVs are computed for each conformation, which is then assigned to a bin (*u, v*) (equations 8-11). For each occupied bin, a representative conformation *f*_*u,v*_ ∈ *S*_*u,v*_ is selected and assigned the total weight of the bin, 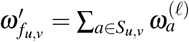. A set of *M* representative conformations 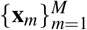is thus obtained and serves as the input ensemble for reweighting against cryo-EM images (equations 13-17).

#### 2. WE simulations with adaptive binning

The reweighted ensemble 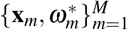serves as starting structures for WE simulations. The WE protocol, including propagation, resampling, and bin assignment in the CV space, is implemented as in equations 3-7. Once the WE simulation is complete, the walker trajectories are combined, and the resulting ensemble is projected onto the CV space (*q*^(1)^, *q*^(2)^).

The weighted probability in each bin 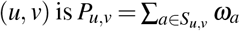 where *S*_*u,v*_ is the is the set of all frames of the combined WE trajectories assigned to bin (*u, v*). The free energy surface obtained from the WE bin probabilities *P*_*u,v*_ is

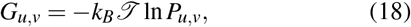

where *k*_*B*_ is the Boltzmann constant and *T* is the temperature. The frame density *ρ*_*u,v*_ of the prior ensemble is the fraction of prior frames assigned to bin (*u, v*). The anchor bins are the occupied bins of highest density within a region *ℛ* of the CV space, 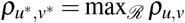. Here, (*u*^∗^, *v*^∗^) are the anchor bin indices in the discretized CV space, and *ℛ* is the tail of the prior distribution along a chosen progress coordinate. The anchor coordinates 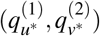 set the outer bin of the binning scheme for the subsequent WE simulation, thereby concentrating the adaptive bins on *R*.

The number of WE subiterations per *cryoWEight* iteration, *N*_WE_, is determined from the effective sample size 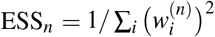 normalized by the walker count 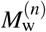, where *n* is the subiteration index. The ratio 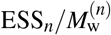 decreases over the first *n*_*d*_ subiterations and varies within a narrow range for *n* > *n*_*d*_, and the cryo-EM reweighting step is performed at *N*_WE_ > *n*_*d*_.

#### 3. Iteration

The reweighting and WE sampling stages are repeated, with each new reweighted ensemble seeding the subsequent simulation. Convergence is assessed from the Kullback-Leibler (KL) divergence between the distributions *P*^(ℓ)^ = {*P*_*u,v*_} of consecutive iterations. The iterations are stopped on ce the divergence falls below a threshold, *D*_KL_(*P*^(*ℓ*+1)^ ∥ *P*^(*ℓ*)^) < *ε*, where *ε* > 0 is a user-defined tolerance. The KL criterion requires no knowledge of the target distribution and therefore remains valid for experimental cryo-EM images. In the current work, the target distribution is known from the synthetic cryo-EM images, and the agreement with the reweighted ensemble is additionally reported as a validation.

### D. System setup

All-atom MD simulations were performed with the OpenMM 8.1^105^ engine in the NVT canonical ensemble with the Amber ff14SB force field^106^. Initial structures were minimized with the limited-memory quasi-Newton algorithm until a tolerance of 10 kJmol^−1^ nm^−1^ or 10,000 iterations was reached.

A Langevin integrator maintained the temperature at 300 K with a time step of 2 fs. For each protein, long-timescale unbiased MD defined the equilibrium distribution from which the synthetic cryo-EM images were generated, and an independent short trajectory initiated from the initial state provided the prior ensemble for reweighting.

The chignolin protein was simulated with the TIP3P explicit water model^107^ in a cubic box with 1.2 nm padding, with particle mesh Ewald (PME) electrostatics^108^ at a cutoff of 1.0 nm and a friction coefficient of 1 ps^−1^. The solvated protein comprised 2,535 atoms, with 10 amino acid residues and 799 water molecules within a cubic box of edge length 3.13 nm. The NTL9 protein was simulated in the OBC2 generalized Born implicit solvent model^109^ with a non-bonded cutoff of 2.0 nm, a solvent dielectric constant of 78.5, and a friction coefficient of 5 ps^−1^, without periodic boundaries, and comprised 627 atoms with 39 amino acid residues. The ADK protein was simulated with the TIP3P explicit water model in a cubic box with 1.2 nm padding, with PME electrostatics at a cutoff of 1.0 nm and a friction coefficient of 1 ps^−1^. The solvated protein comprised 31,137 atoms, with 214 amino acid residues, 9,264 water molecules, and four sodium counterions within a cubic box of edge length 6.92 nm. Each *cryoWEight* iteration comprised 30 WE subiterations at *τ* = 20 ps for chignolin, 40 subiterations at *τ* = 40 ps for NTL9, and 25 subiterations at *τ* = 100 ps for ADK (Supplementary Tables S1-S3).

WE simulations were performed with the WESTPA 2.0 software^110^ with a target of four walkers per bin. For trajectory propagation, WESTPA 2.0 employs the OpenMM 8.1 engine, with the force field, solvent model, and integrator settings for each protein retained from the MD simulation that generated the prior ensemble. The CV space was discretized employing the rectilinear grid and the MAB scheme, as described in Section II A.

#### 1. Chignolin

Chignolin is a synthetic 10-residue mini-protein that adopts a stable *β* -hairpin fold in aqueous solution^111,112^. Despite its small size, chignolin exhibits folding and unfolding behavior with a well-defined native structure stabilized by intramolecular hydrogen bonds and aromatic interactions^27,113,114^. The microsecond timescale of chignolin folding is computationally achievable, and with well-characterized thermodynamics and kinetics, it serves as a benchmark system for validating enhanced sampling methods^39,115–117^, machine learning approaches^118–121^, and force field development^106,122,123^. The initial protein structure was obtained from the Protein Data Bank (PDB ID: 1UAO)^111^. Long-timescale MD simulations were performed to define the reference equilibrium distribution from which the synthetic cryo-EM images were generated. Equilibrium sampling comprised four 20 µs trajectories initiated from the folded state and four 15 µs trajectories initiated from the unfolded state, for an aggregate simulation time of 140 µs across 1.4 × 10^7^ frames (Supplementary Table S1a). CVs were defined as the root mean square deviation (RMSD) and the radius of gyration (*R*_*g*_), both computed relative to the folded state of chignolin. Frames with RMSD ≥ 7.25 Å were classified as unfolded, and 10,000 frames were sampled from this subset to form the target ensemble for image generation (Supplementary Table S1b). A 200 ns MD simulation initiated from the folded state generated the starting conformations for reweighting. The reweighted conformations then seeded the WE simulations with *cryoWEight* (Supplementary Tables S1c and S1d).

#### 2. NTL9

The N-terminal domain of ribosomal protein L9 (NTL9) is a 39-residue protein that adopts a three-stranded anti-parallel *β* -sheet packed against a single *α*-helix^124,125^. The protein folds reversibly via a two-state mechanism with a folding time of approximately 1.5 ms^125,126^, thereby establishing it as a benchmark system for enhanced sampling methods^38,39,41^ and protein folding studies^126–129^. The reference ensemble was constructed from eight independent trajectories with four simulation replicates of length 25 µs propagated from the NTL9 folded state and four simulations of length 20 µs propagated from the unfolded state, spanning 180 µs across 1.8 × 10^7^ frames (Supplementary Table S2a). CVs were defined as RMSD and *R*_*g*_ relative to the folded reference state of NTL9.

Frames with RMSD ∈ [2.00, 3.00] Å were classified as folded, and 10,000 frames were sampled from this subset to form the target ensemble for image generation (Supplementary Table S2b). A 200 ns MD simulation initiated from the unfolded state generated the starting conformations for reweighting, which then seeded the WE simulations (Supplementary Tables S2c and S2d).

#### 3. Adenylate kinase

Among the nucleoside monophosphate kinase superfamily, adenylate kinase (ADK) is a 214-residue enzyme that catalyzes the reversible phosphoryl transfer reaction ATP + AMP ⇌ 2 ADP, central to cellular energy homeostasis^130^.

ADK displays a well-characterized large-scale conformational transition between ligand-free open and substrate-bound closed states^131–135^. The ADK enzyme comprises three principal domains, i.e., a relatively rigid CORE domain, an adenosine monophosphate (AMP)-binding nucleoside monophosphate (NMP) domain, and an adenosine triphosphate (ATP)-binding LID domain^136^. The NMP and LID domains undergo large-scale hinge-bending motions relative to the CORE domain, which are well characterized using angular order parameters^131,137^. Long-timescale simulations were initiated from the closed conformational state of ADK (PDB ID: 1AKE)^136^, with four independent simulation runs of length 9 µs each with 3.0 × 10^5^ frames, yielding an aggregate simulation time of 36 µs across 1.2 × 10^6^ frames for the reference equilibrium distribution (Supplementary Table S3a). CVs were defined for each frame as the opening angles of the NMP and LID domains relative to the CORE domain (*θ*_NMP_ and *θ*_LID_). Frames with *θ*_NMP_ ∈ [70^°^, 85^°^] and *θ*_LID_ ∈ [130^°^, 150^°^] were classified as open states, and 2,500 frames were sampled from this subset to form the target ensemble for image generation (Supplementary Table S3b). The prior ensemble for reweighting was drawn from a 20 ns MD simulation of the closed state (Supplementary Table S3c).

## III. Results

WE simulations and cryo-EM reweighting establish a directional iterative loop in which each reweighting cycle progressively drives the simulation ensemble towards the cryo-EM target distribution. The convergence towards the target distribution primarily depends on two factors, i.e., (i) the information content of the cryo-EM images, governed by the SNR, and (ii) the WE path sampling strategy, governed by parameters such as the resampling time, binning scheme, and the number of walkers per bin. The following sections describe the convergence onto a non-image target (Section III A) and onto the cryo-EM target (Section III B), the binning dependence (Section III C), the signal strength dependence (Section III D), the multimodal recovery (Section III E), and the functional transition pathways (Section III F).

### A. Iterative reweighting converges onto a non-image target distribution

The ensemble reweighting algorithm updates the conformational weights only through the per-structure likelihood, and the iterative protocol is therefore first evaluated as a proof of concept on a target distribution defined independently of cryo-EM particle images. The iterative reweighting was examined on the chignolin protein in the OBC2 generalized Born implicit solvent model^109^, with a non-bonded cutoff of 2.0 nm, a solvent dielectric constant of 78.5, and a friction coefficient of 1 ps^−1^. The target distribution was defined in the CV space by the root mean square deviation (RMSD) ∈ [4, 6] Å. A 200 ns unbiased MD simulation of folded chignolin provided the initial ensemble at C*α*-RMSD ∈ [0, 2] Å, with the per-structure likelihood constructed from the distance between each conformation and the target distribution in the CV space under a Gaussian kernel of bandwidth 0.6 Å. Each iteration comprised 15 WE subiterations at *τ* = 20 ps, with a target of four walkers per bin. Over successive iterations of WE sampling and reweighting, the ensemble converged onto the target distribution (Figure 2b), with the sampled conformations partially unfolded rather than fully extended (Figure 2a). Once the ensemble reached the target distribution, it remained centered across all subsequent iterations, fluctuating within the target band without repopulating the folded basin at lower RMSD values. An unbiased WE simulation instead recovers the equilibrium distribution, and the higher-RMSD target is sampled only when the ensemble is reweighted to match the external signal.

**FIG. 2:**
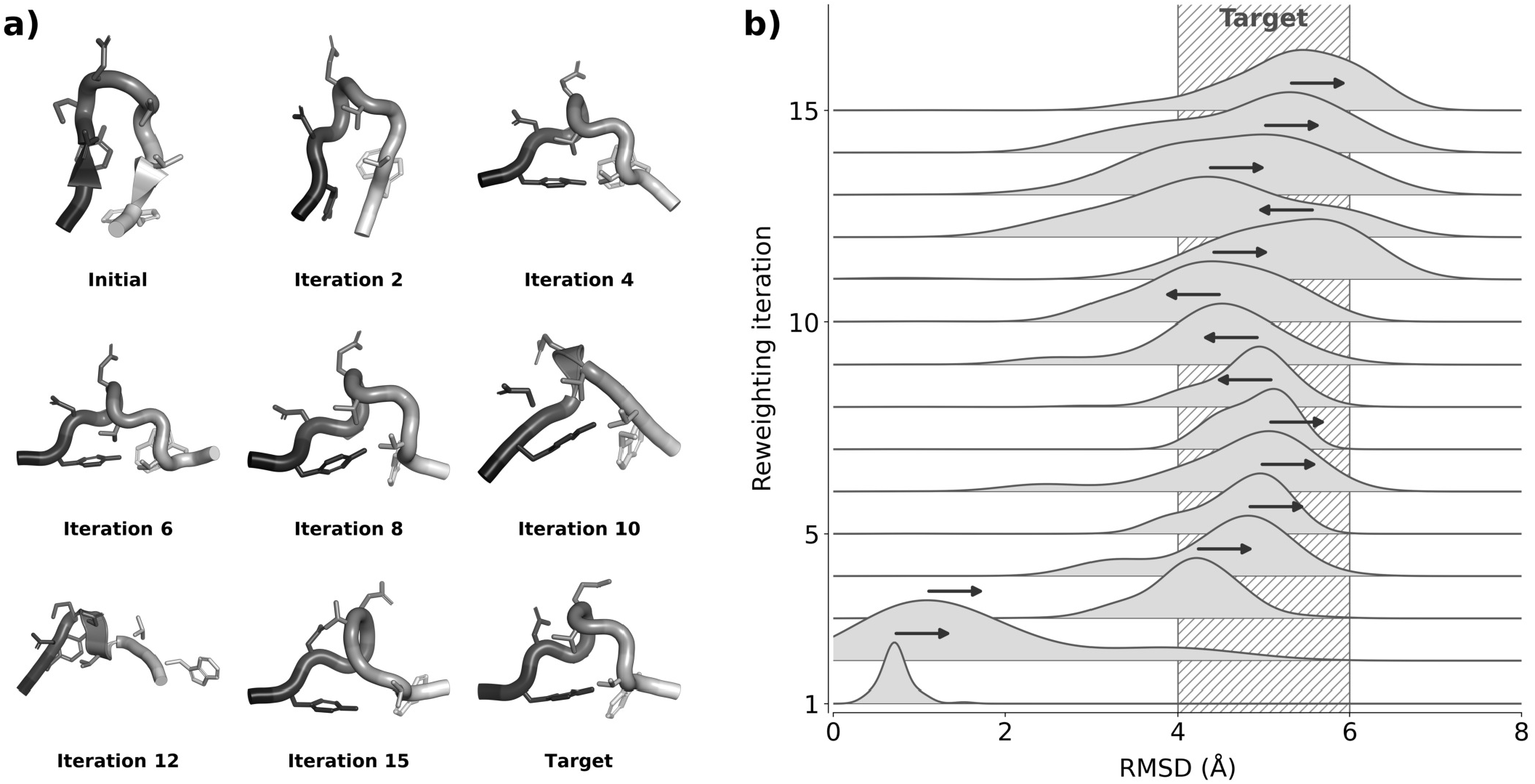
Iterative reweighting protocol for the chignolin protein in implicit solvent with a target distribution defined by root mean square deviation (RMSD) ∈ [4, 6] Å. (a) Representative conformations are shown for the initial state, at iterations 2, 4, 6, 8, 10, 12, and 15, and for the target distribution. (b) The RMSD distribution for each reweighting iteration is plotted, and the arrows indicate the direction of movement between successive iterations. Starting from the folded state, the distribution converges onto the target over successive iterations.

### B. *cryoWEight* converges onto target distributions

*cryoWEight* was first evaluated on the chignolin protein against cryo-EM images at SNR = 1.0 (Figure 3a), a high-signal regime where images contain strong discriminatory information. The initial ensemble is systematically driven along RMSD and *R*_*g*_ from the folded basin to the non-overlapping cryo-EM distribution (Figure 3a). Over four successive iterations, conformations evolve from compact *β* -hairpin folds through partially folded intermediates towards extended unfolded states that overlap with the cryo-EM populations, with the free energy landscape progressively moving from the folded to the unfolded region over a total sampling time of 1.48 µs (Figure 3a). The posterior weight densities redistribute from low to high RMSD and *R*_*g*_ (Figure 3b), as cryo-EM reweighting assigns higher weights to conformations whose projections most closely match the images. The redistributed weights then propagate to the next WE resampling step, where walkers in overpopulated bins are merged, and walkers in sparsely populated bins are split, thereby amplifying sampling in the target region. Distribution dissimilarity metrics, i.e., the Kullback-Leibler (KL) divergence, the Jensen-Shannon (JS) divergence, the Bhattacharyya distance, and the Wasserstein distance^138^, decreased monotonically across iterations relative to the cryo-EM target (Figure 3c and Supplementary Table S4). The statistical quality of the ensembles is further evaluated employing the effective sample size 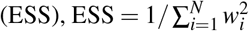 where {*w*_*i*_} are the normalized frame weights^139^. When all weights are uniform (*w*_*i*_ = 1/*N*), ESS equals *N* and all frames contribute equally, whereas low ESS values indicate that the probability mass has concentrated on a few structures, with the remainder carrying vanishing weights. Across iterations, the ESS/*N* ratios ranged from 3-11% (Figure 3c), indicating effective discrimination between conformations while maintaining sufficient ensemble diversity to avoid collapse (Supplementary Table S5).

**FIG. 3:**
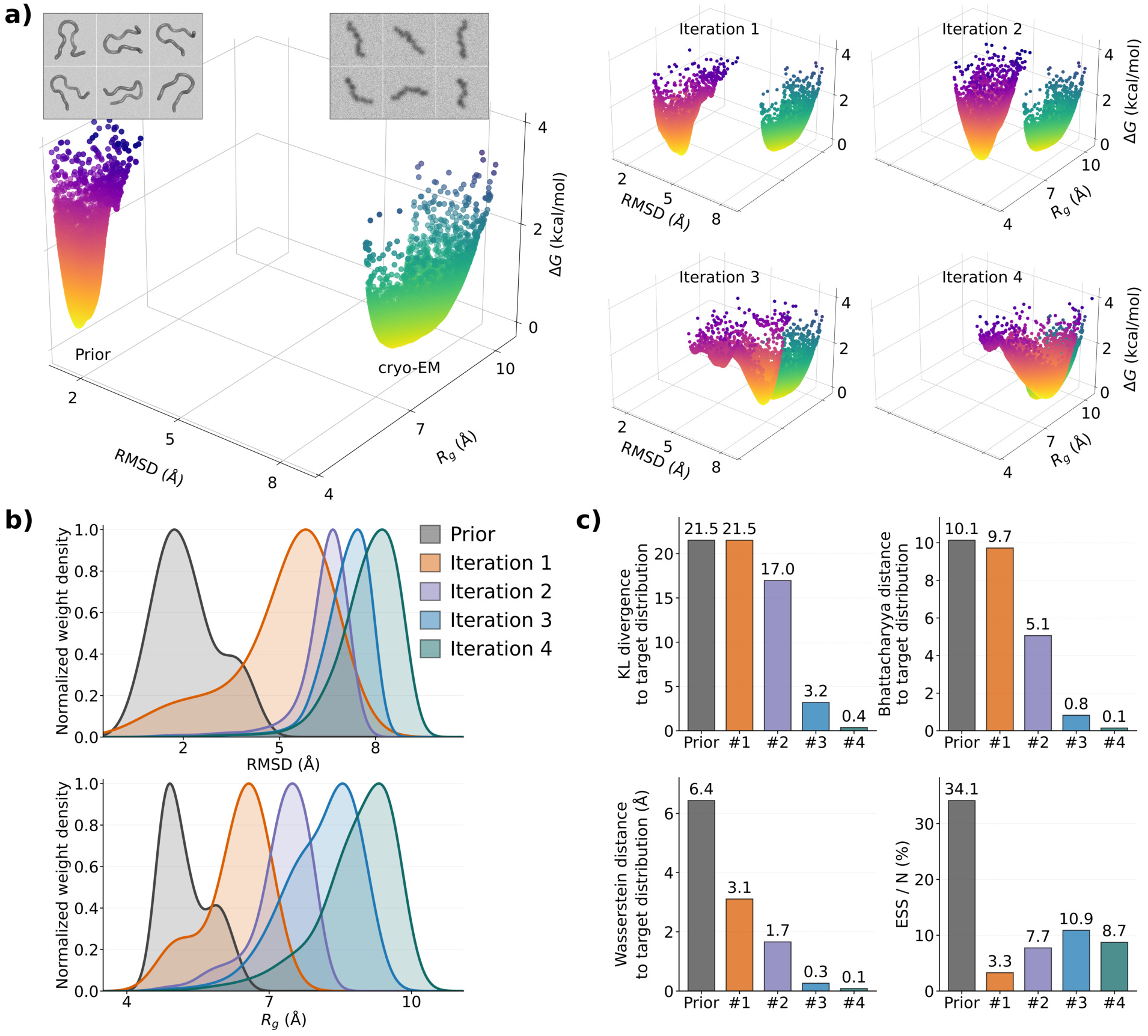
*cryoWEight* implemented to chignolin protein with cryo-EM images at SNR = 1.0. (a) Initial conformational ensemble of folded chignolin from a 200 ns molecular dynamics simulation and the cryo-EM ensemble of the unfolded state, both as 3D scatter plots over RMSD, *R*_*g*_, and free energy, with representative prior conformations and synthetic cryo-EM particle images above the corresponding ensemble. Free energy landscapes are shown against the cryo-EM ensemble for four successive *cryoWEight* iterations (right) as the ensemble transitions from the folded basin to the cryo-EM distribution. (b) Posterior weight density along RMSD (top) and *R*_*g*_ (bottom) for the prior and four successive iterations. (c) Distribution dissimilarity metrics against the cryo-EM target across iterations, i.e., Kullback-Leibler divergence, Bhattacharyya distance, and Wasserstein distance, all decrease monotonically. The fourth subplot shows the effective sample size ratio ESS/*N* (%) for the prior and each iteration.

*cryoWEight* was next implemented to the NTL9 protein in implicit solvent, with the initial unfolded ensemble reweighted against folded cryo-EM images at RMSD ∈ [2, 3] Å and SNR = 1.0 (Figure 4a). Over four iterations and a sampling time of 6.45 µs, the free energy landscapes show the global minimum relocating from high to low RMSD and *R*_*g*_ values, consistent with the unfolded-to-folded transition (Figure 4a). To further demonstrate the convergence of the current protocol to arbitrary target distributions rather than a specific conformational state, an alternative cryo-EM target distribution was defined with a narrower RMSD range of 2.75 Å-3.50 Å for the folded NTL9 states. Despite a different target definition, *cryoWEight* simulations successfully converged within three iterations and a sampling time of 4.58 µs (Supplementary Table S6). Sampled conformations evolved from extended unfolded states through partially folded intermediates to the native NTL9 fold (Figure 4b), and the posterior weight density along RMSD redistributed from high to low values across iterations (Figure 4c). The narrower target followed the same progression. The distribution dissimilarity metrics decreased with respect to the cryo-EM target distribution across iterations in both cases (Supplementary Table S6).

**FIG. 4:**
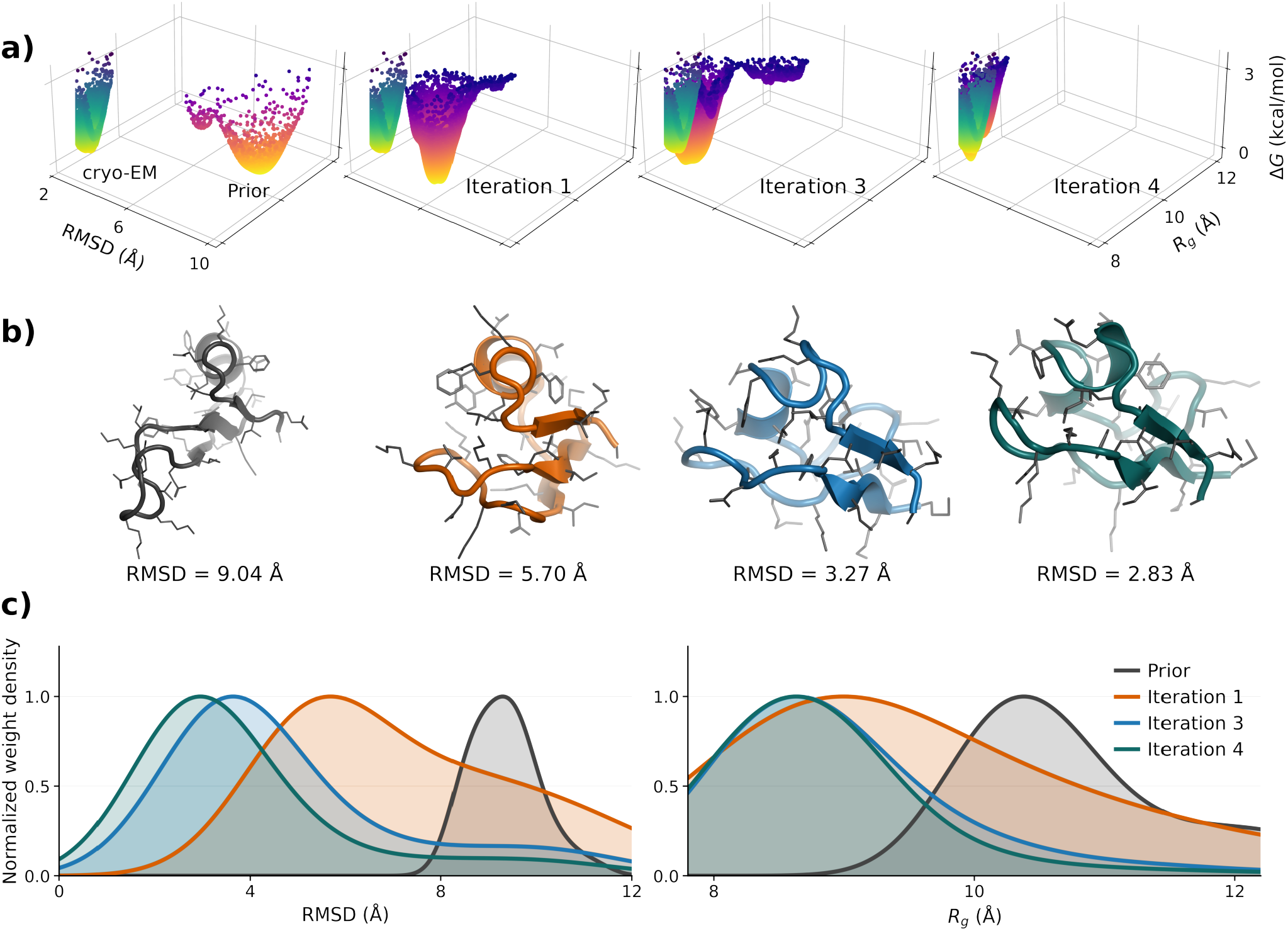
*cryoWEight* implemented to the NTL9 protein with cryo-EM images at SNR = 1.0. (a) The initial conformational ensemble from a 200 ns molecular dynamics simulation and the cryo-EM ensemble of folded conformations defined by RMSD ∈ [2, 3] Å are shown as 3D scatter plots over RMSD, *R*_*g*_, and free energy. Free energy landscapes at successive *cryoWEight* iterations (right) show the transition of the ensemble from the unfolded basin to the cryo-EM distribution. (b) Representative NTL9 structures are shown for the prior and iterations 1, 3, and 4, from the extended unfolded conformation through partially folded intermediates to the native fold conformation. The RMSD to the folded reference is given below each structure. (c) Posterior weight density along RMSD (left) and *R*_*g*_ (right) for the prior and successive iterations.

*cryoWEight* was then assessed on the ADK protein, with an initial closed ensemble reweighted against cryo-EM images of the open state defined by *θ*_NMP_ ∈ [70^°^, 85^°^] and *θ*_LID_ ∈ [130^°^, 150^°^] at SNR = 1.0. Over seven iterations and a sampling time of 13.26 µs, conformations evolved from the compact closed state, with the NMP and LID domains positioned close to the CORE domain, through partially open intermediates, to extended open conformations with both domains rotated outward (Figure 5b). This progression captures the well-characterized hinge-bending motions of the NMP and LID domains relative to the rigid CORE domain, spanning the conformational coordinate between the open and closed catalytic states of ADK. CV distributions along *θ*_NMP_ transition from a narrow distribution centered around 45^°^ in the prior through broader transitional intermediate distributions to a distribution centered around 75^°^ at convergence. Along *θ*_LID_, the distribution broadens substantially from a narrow prior around 115^°^ to a broad distribution spanning 90^°^ to 180^°^ in later iterations, indicating a larger range of LID conformations accessible in the open state (Figure 5a). The posterior weight density along *θ*_NMP_ redistributes from low to high angular values, with the initial ensemble concentrated below 60^°^ and successive iterations shifting onto the cryo-EM range of [70^°^, 85^°^]. The posterior weight density along *θ*_LID_ similarly shifts from the prior around 115^°^ towards 150^°^ across iterations, converging onto the cryo-EM target range of [130^°^, 150^°^] (Figure 5a). The distribution dissimilarity metrics decrease substantially over successive iterations but plateau at non-zero values rather than approaching zero (Supplementary Table S7). This behavior arises from the larger system size of ADK, where conformational heterogeneity orthogonal to *θ*_NMP_ and *θ*_LID_ persists at convergence and contributes to residual differences relative to the cryo-EM target. The ESS/*N* ratios ranged from 2-6% across iterations, indicating effective discrimination between conformations and sufficient ensemble diversity to avoid weight collapse (Supplementary Table S8).

**FIG. 5:**
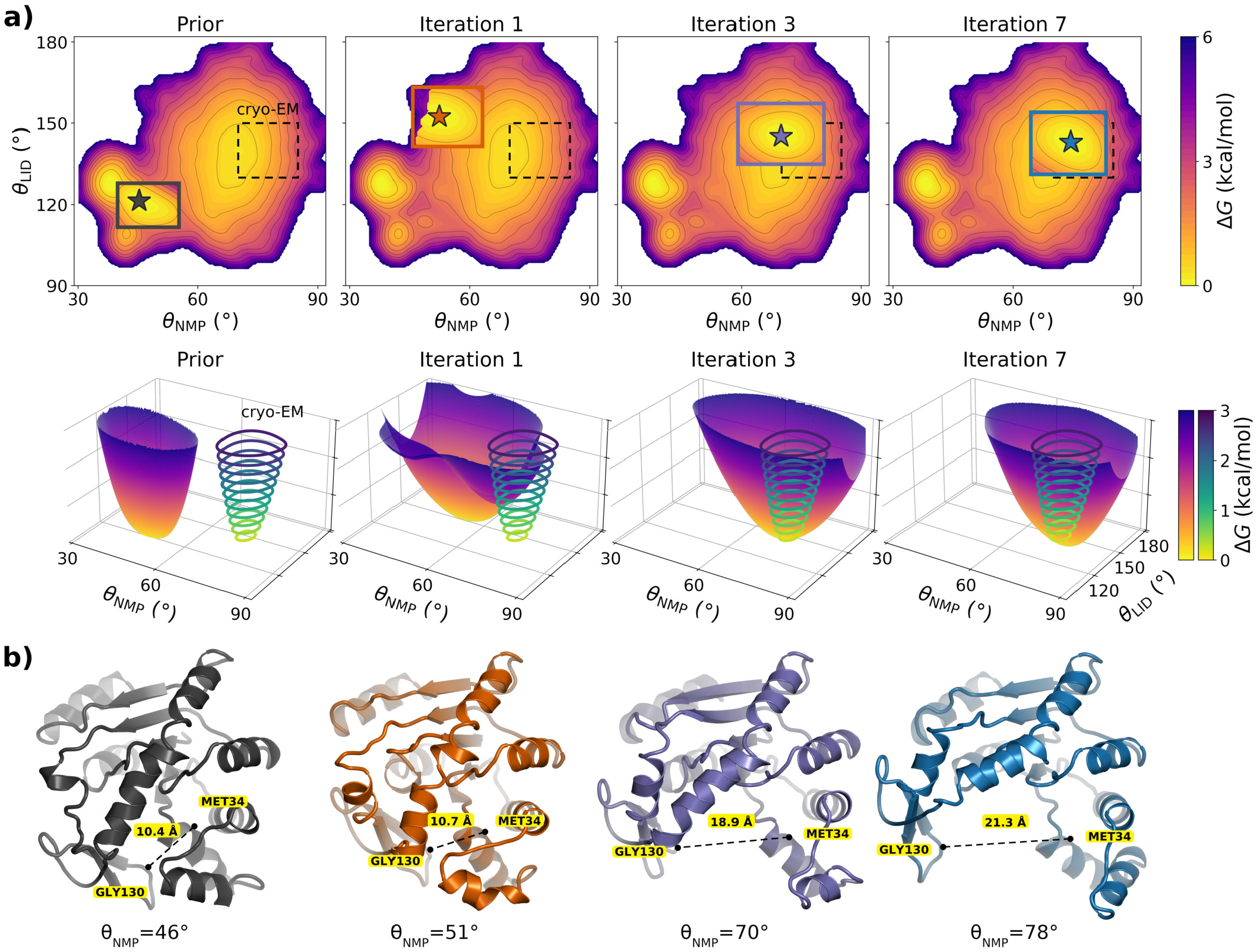
*cryoWEight* implemented to the adenylate kinase (ADK) protein with cryo-EM target distribution at SNR = 1.0. (a) Free energy landscapes over *θ*_NMP_ and *θ*_LID_ are shown for the prior and successive *cryoWEight* iterations. The top row shows the reference free energy landscape from 36 µs of unbiased molecular dynamics simulations in each panel, and the *cryoWEight* ensemble free energy is drawn within the basin of its free energy minimum. The cryo-EM target distribution spans *θ*_NMP_ ∈ [70^°^, 85^°^] and *θ*_LID_ ∈ [130^°^, 150^°^]. The bottom row shows the free energy surface of the reweighted *cryoWEight* ensemble for each iteration, with the cryo-EM target free energy shown as free energy contours. (b) Representative ADK conformations at each iteration, from the closed state through partially open intermediates to the fully open state. The C*α* separation between Met34_NMP_ and Gly130_LID_ is given for each structure.

### C. *cryoWEight* remains invariant to binning

A WE simulation is sensitive to the conformational space partition through the target number of walkers per bin and the location of the adaptive bins. The contribution of the binning strategy to the recovered ensemble is therefore distinguished from that of the reweighting. In each *cryoWEight* iteration, the walker weights are corrected by cryo-EM reweighting, and the anchor that sets the outer bin location is recomputed from the prior trajectory. The anchor is defined as the occupied bin of highest frame density beyond the cutoff *µ* + *kσ*, where *µ* and *σ* are the mean and standard deviation of the prior along the progress coordinate *ξ*, and *k* is the number of standard deviations. Successive values |*k* |= 3, 2, and 1 denote the distal, medial, and proximal anchors and are evaluated until occupied bins remain beyond the cutoff (Figure 6a).

**FIG. 6:**
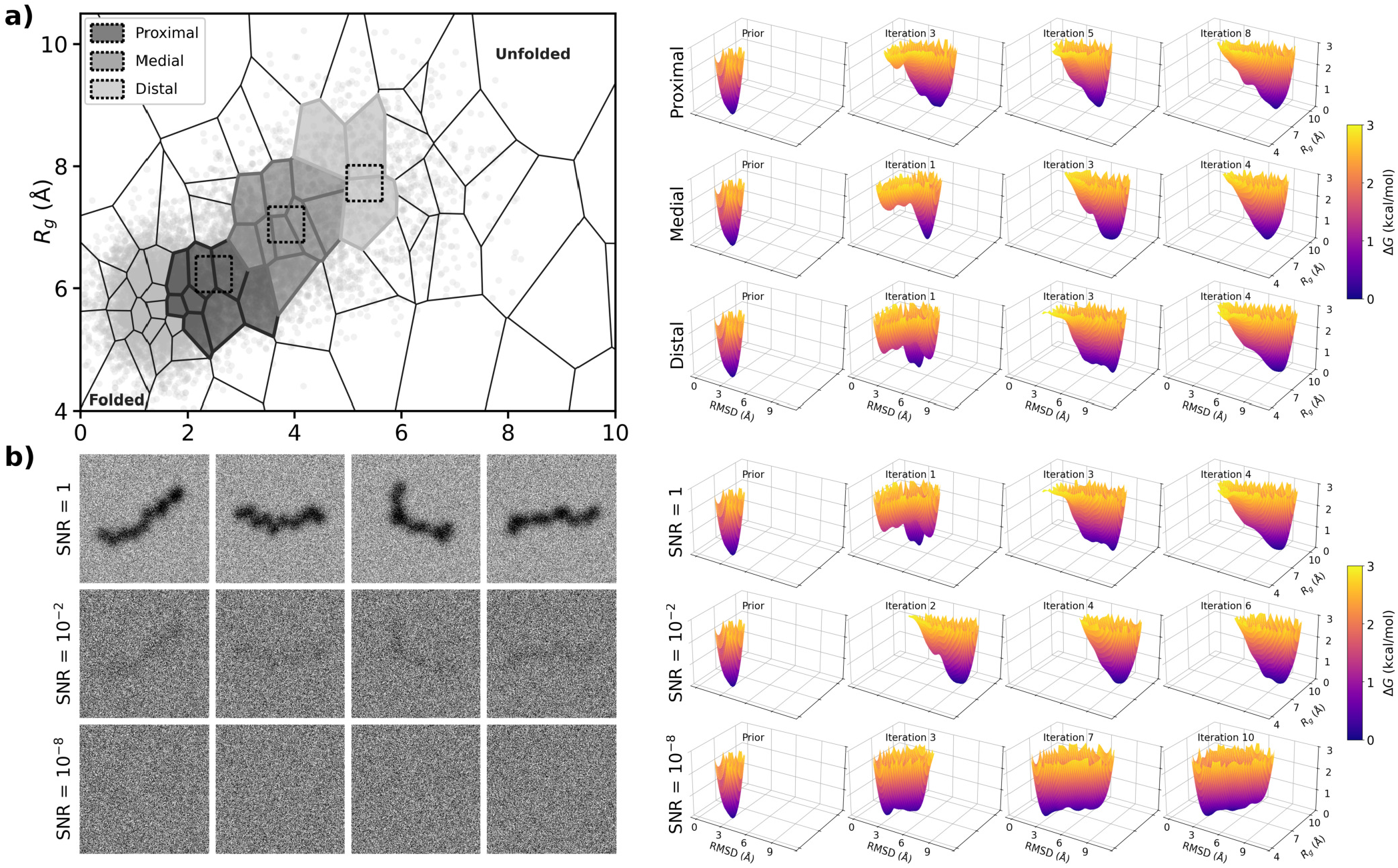
Dependence of *cryoWEight* on the minimal adaptive binning (MAB) anchor location within the weighted ensemble (WE) simulations and on the cryo-EM SNR for the chignolin protein. (a) The MAB anchor is varied, and the ensemble is iteratively reweighted against cryo-EM images at SNR = 1.0. The RMSD-*R*_*g*_ space is partitioned into Voronoi cells, and the distal, medial, and proximal anchors, marked by dashed boxes within the shaded cells, are obtained at cutoffs |*k*| = 3, 2, and 1 along the progress coordinate between the folded and unfolded basins of chignolin (left). The free energy landscapes as functions of RMSD and *R*_*g*_ are shown for each anchor across the *cryoWEight* iterations (right). (b) The cryo-EM SNR is varied for the distal MAB anchor. Representative cryo-EM particle images correspond to SNR = 1.0, 10^−2^, and 10^−8^ (left). The free energy landscapes as functions of RMSD and *R*_*g*_ are shown for each SNR across the *cryoWEight* iterations (right), where convergence to the cryo-EM target distribution fails only at SNR = 10^−8^.

The dependence of the recovered ensemble on the anchor location was assessed for the chignolin protein, with the anchor placed at three different locations along the progress coordinate, namely distal, medial, and proximal (Figure 6a). The proximal anchor sits within the densest region of the prior CV distribution, the distal anchor in a sparsely populated region farther from the prior basin, and the medial anchor between the two. All three anchor locations converged to the same cryo-EM distribution, establishing the binning-invariance of the iterative reweighting protocol and the independence of the final ensemble from the choice of MAB anchor. The distal anchor reached the cryo-EM distribution within four iterations and a sampling time of 1.48 µs, the medial anchor also in four iterations but at 1.82 µs, and the proximal anchor in eight iterations and 3.38 µs (Figure 6a), reflecting the additional WE sampling required to populate the sparsely sampled high-*ξ* region farther from the prior basin.

### D. *cryoWEight* depends on cryo-EM signal strength

Having established convergence at SNR = 1.0, the dependence of *cryoWEight* on the cryo-EM signal strength was next evaluated for chignolin by lowering the SNR. At SNR = 10^−2^, the ensemble converged within six iterations with a sampling time of 2.79 µs (Figure 6b). At SNR = 10^−8^, the image signal is buried in the noise and the protocol failed to converge even after 3.38 µs of sampling (Figure 6b). Consistent with this failure, the effective sample size ratio ESS/*N* approached nearly 100% at SNR = 10^−8^, indicating nearly uniform weights across all frames and confirming the absence of any discriminatory information from the cryo-EM signal (Supplementary Table S5). The failure at SNR = 10^−8^ also establishes that convergence at higher SNR is driven by the cryo-EM signal rather than by an intrinsic bias of the WE sampling protocol.

The signal dependence was reproduced for NTL9 protein, where the reweighting protocol failed to converge at SNR = 10^−8^ with no directional shift towards the target distribution, even after 15.76 µs of sampling. For converged simulations at SNR = 1.0, the ESS/*N* ratios ranged from 2-12%, indicating effective discrimination between conformations, whereas at SNR = 10^−8^ the ESS/*N* ratio remained near 100% throughout all iterations, confirming the absence of any discriminatory information in the cryo-EM signal (Supplementary Table S9).

### E. *cryoWEight* recovers multimodal populations at controlled ratios

Beyond convergence to a single target state, *cryoWEight* recovered multimodal target distributions of coexisting conformational states at controlled relative populations for chignolin protein. Initiating from the folded state, conventional WE simulations were run for 50 subiterations, reweighted against cryo-EM images sampled from a mixture of folded and unfolded chignolin conformations in multiple ratios, and propagated for 50 additional WE subiterations from the reweighted ensemble. Conformations were then classified into three states, i.e., folded (RMSD < 2 Å), intermediate (2 Å ≤ RMSD ≤ 6 Å), and unfolded (RMSD > 6 Å). The recovered *cryoWEight* RMSD distributions track the cryo-EM target distributions across the folded to unfolded ratios (Figure 7a). Across the nine folded to unfolded cryo-EM population ratios, the recovered folded population in the *cryoWEight* ensemble increases monotonically from 10% to 86% as the cryo-EM folded fraction increases from 10% to 90% (Figure 7b), agreeing quantitatively with the cryo-EM target with Spearman *ρ* = 1.00 (Figure 7c). The recovered unfolded population follows the cryo-EM target with *ρ* = 0.85 (Figure 7c) but lies consistently below it. The shift is most pronounced at high unfolded populations, where the recovered intermediate states exceed the recovered unfolded population (Figure 7b). Although cryo-EM reweighting assigns higher weights to walkers in unfolded conformations, the unbiased dynamics propagated during the subsequent WE subiterations favor the folded state at thermodynamic equilibrium, redistributing the ensemble towards folded conformations. Despite this redistribution, the monotonic recovery demonstrates the directional bias of *cryoWEight* towards unfolded conformations.

**FIG. 7:**
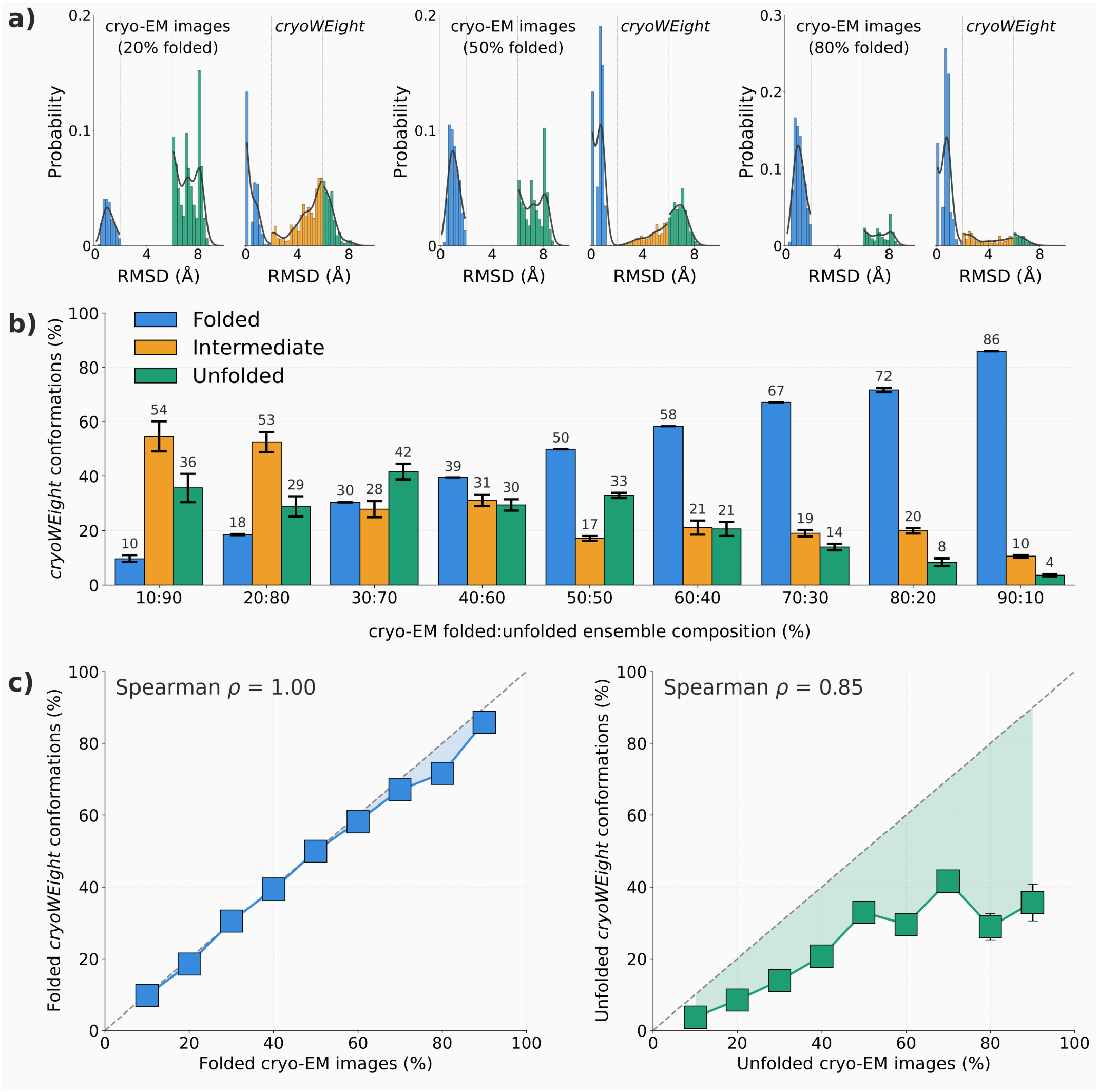
Multimodal reweighting of chignolin with *cryoWEight*. Starting from the folded state, weighted ensemble (WE) simulations are run for 50 subiterations, reweighted against cryo-EM images sampled at different folded-to-unfolded ratios ranging from 10:90 to 90:10, and propagated for a further 50 subiterations from the reweighted ensemble. (a) The cryo-EM and *cryoWEight* RMSD distributions are compared at three of these ratios (20:80, 50:50, and 80:20). Conformations below 2 Å C*α* RMSD are folded (blue), those between 2 Å and 6 Å are intermediate (orange), and those above 6 Å are unfolded (green). (b) The recovered populations of the folded, intermediate, and unfolded conformations are shown for all nine ratios. (c) The recovered folded population is plotted against the folded fraction in the cryo-EM images (left), and the recovered unfolded population is plotted against the unfolded fraction (right), with Spearman rank correlations of *ρ* = 1.00 and *ρ* = 0.85, respectively. Error bars are the standard deviation over the last five WE subiterations.

### F. *cryoWEight* recovers the functional transition pathway

Long timescale unbiased MD simulations, which record multiple transitions between the closed and open states over 36 µs of aggregate sampling across four replicas, serve as the reference functional pathway against which the *cryoWEight* pathway is validated. To order configurations along the transition, the committor *φ* (x) ∈ [0, 1] is computed from a Markov state model obtained from the unbiased MD simulations over the *θ*_NMP_-*θ*_LID_ space. The *θ*_NMP_-*θ*_LID_ CV space is partitioned into 150 microstates by k-means clustering, and a transition probability matrix T is estimated from the microstate transitions observed at a lag time of 2.4 ns^140^. The committor is the harmonic solution *φ*_*i*_ = ∑ _*j*_ *T*_*i j*_ *φ*_*j*_ of the transition matrix ^141^ over the intermediate states, with the boundary conditions *φ*_*i*_ = 0 in the closed macrostate and *φ*_*i*_ = 1 in the open macrostate, where the macrostates are defined by the LID angle as *θ*_LID_ ≤ 110^°^ for the closed state and *θ*_LID_ ≥ 145^°^ for the open state. Each configuration in both the reference MD and the *cryoWEight* ensemble is assigned the committor of its nearest microstate, which places it along the closed to open transition. The committor serves solely as a progress coordinate that orders the transition, with no kinetic quantities inferred from the model, and the transition matrix is used only to define this ordering and not to extract rates.

The interdomain motions of the *cryoWEight* ensemble are projected onto the committor and compared against the reference MD pathway (Figure 8). Two interdomain contacts describe the opening. The Arg131_LID_-Arg71_CORE_ distance tracks the LID domain relative to the CORE domain (Figure 8a), and the Met34_NMP_-Gly130_LID_ distance tracks the NMP domain relative to the LID domain (Figure 8b). Along both contacts, the reference pathway deviates from the straight interpolation between the closed and open states, a nonlinear transition in which the NMP and LID domains rearrange asynchronously^131,132,134^. Over the committor range, the *cryoWEight* pathway follows the reference pathway to within .5 Å on average, and the representative structures at committor = 0.40 and 0.80 show the same intermediate geometries along the transition (Figure 8 (i-iv)). Because *cryoWEight* initiates from the closed ensemble and is guided only by the cryo-EM images in the open state, the agreement with the reference pathway shows that the external signal shifts the populations along the transition without displacing the intermediates from the unbiased pathway. The straight interpolation is included as a measure of nonlinearity rather than as an outcome of the dynamics (Figures 8a and 8b).

**FIG. 8:**
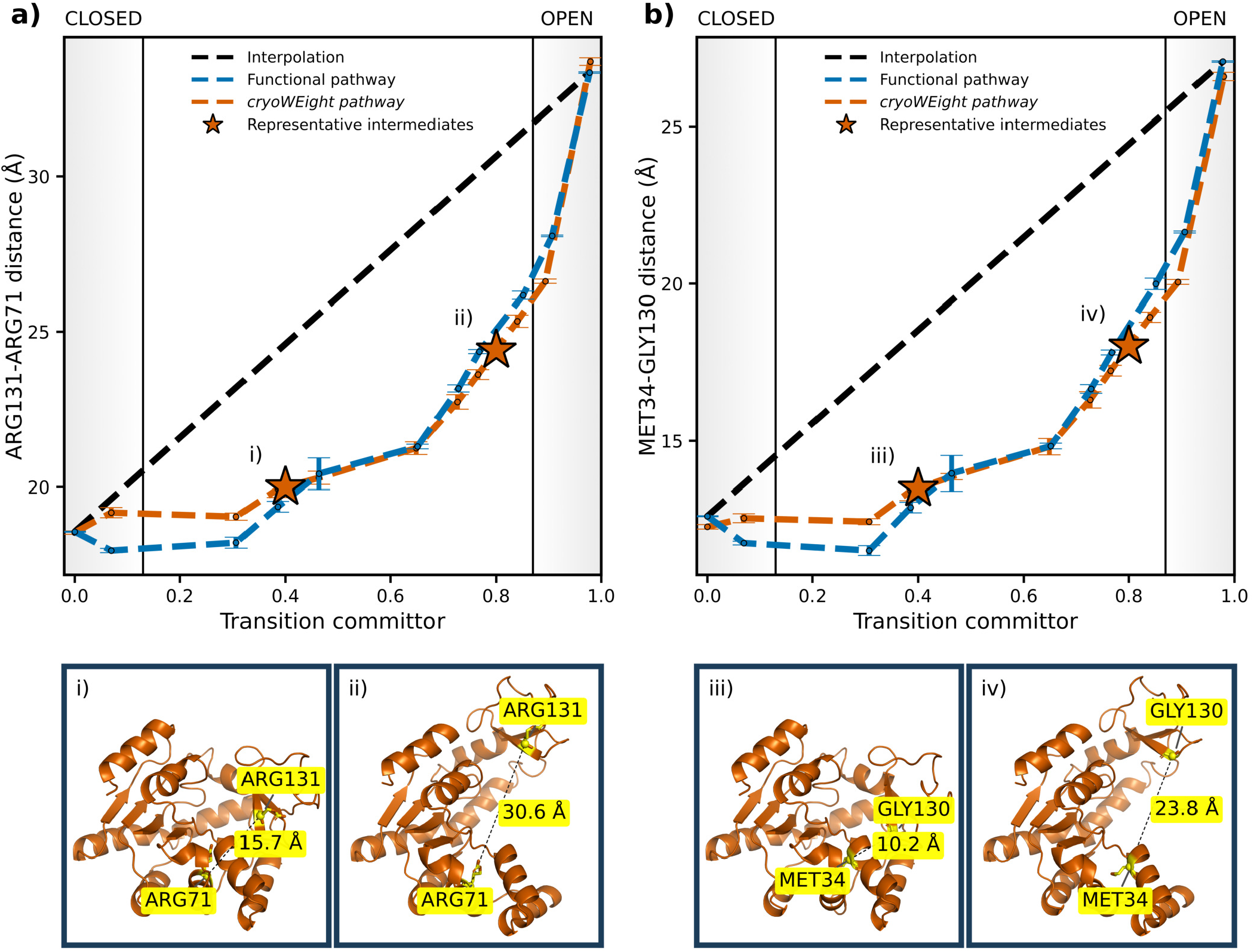
*cryoWEight* recovers the functional transition pathway of ADK between the closed and open states. (a, b) Interdomain C*α*-C*α* distance as a function of the transition committor from the closed to the open state for (a) the Arg131_LID_-Arg71_CORE_ contact and (b) the Met34_NMP_-Gly130_LID_ contact. Each panel compares the functional pathway from unbiased molecular dynamics (MD) simulations (blue), the *cryoWEight* pathway (orange), and the linear interpolation between the endpoints of the functional pathway (black). The shaded regions at either end are the closed and open basins below committor 0.13 and above 0.87. Error bars denote the standard error of the mean over configurations within each committor bin. Markers i-iv lie on the *cryoWEight* pathway at the committor positions of the representative structures shown below. (i-iv) Representative *cryoWEight* conformations of ADK, with panels (i, ii) displaying the Arg131_LID_-Arg71_CORE_ contact at committor = 0.40 and 0.80, respectively, and panels (iii, iv) the Met34_NMP_-Gly130_LID_ contact at committor = 0.40 and 0.80, respectively. The C*α* separation labeled in each conformation is that of the individual structure rather than the binned average plotted above. The separation of the NMP and LID domains is nonlinear, and *cryoWEight* reproduces the functional pathway to within 0.5 Å on average.

## IV. Discussion and Conclusion

*cryoWEight* couples WE sampling with cryo-EM ensemble reweighting because posterior reweighting alone cannot generate conformations missing from the prior trajectory ensemble, and WE simulations alone lack an external reference to direct sampling toward the equilibrium distribution. The framework is validated on three proteins of increasing complexity and in each case, initial free energy landscapes progressively converged towards cryo-EM distributions over successive iterations. Target distributions were recovered across different MAB anchor locations and SNR ratios, failing only when the image signal was buried in noise. For the chignolin protein, the ensemble was driven from the folded basin onto the cryo-EM distribution of the unfolded state, and relative populations were recovered across nine folded-to-unfolded mixtures, with the unfolded fraction underestimated when it dominated the images. For the NTL9 protein, the unfolded ensemble converged onto the folded state with two different target definitions. For the ADK protein, the closed conformational ensemble populated the open state, and the recovered transition followed the pathway obtained from long timescale unbiased MD. In conventional WE simulations, walker weights decrease across successive resampling cycles, leading to weight degeneracy and inefficient exploration of the conformational space. Within *cryoWEight*, walkers are iteratively reweighted against cryo-EM images and reassigned weights based on their likelihood of generating the observed images, thereby facilitating walkers to cross free energy barriers and to populate previously inaccessible regions.

Conventional importance reweighting in the configuration space may fail when the prior ensemble assigns zero probability to conformations favored by the target distribution. The per-structure likelihood remains strictly positive at any finite noise level such that no conformation is assigned zero weight. The expectation-maximization step redistributes weights towards higher likelihood conformations, and subsequent WE sampling generates conformations not represented in the prior ensemble. Neither step requires the prior and the target to overlap in the conformational space. Convergence without overlap is therefore governed by the likelihood itself rather than by the observable that defines it, and the same behavior is obtained for a target specified by a progress coordinate with no particle images (Section III A and Figure 2b). The observable instead determines the discrimination resolution, and a likelihood defined on the progress coordinate cannot distinguish conformations degenerate along it, whereas the image likelihood separates such conformations through their 2D projections. Any observable that defines a per-structure likelihood can serve as the target, and conformational distributions from NMR^142–144^ and SAXS^145,146^ can therefore inform the weights of the resulting ensembles. When the prior ensemble populates the target states, reweighting alone recovers their populations^77^, and without such an overlap, the target conformations must be generated by the sampling rather than recovered by reweighting alone. The absence of overlap between the prior and the target was therefore imposed by design in all three proteins.

The WE methodology depends on predefined progress coordinates to bin walkers for splitting and merging operations, a limitation shared by metadynamics^102,147–149^, milestoning^150–153^, and related enhanced sampling methods^23,154,155^. In each case, the coordinate is specified in advance rather than obtained from the experimental data. Conformational coordinates have recently been derived directly from particle images with methods such as manifold embedding and neural latent spaces^61–63^. An image-derived coordinate is likewise available from the structure-image log likelihood matrix evaluated in the reweighting step. Its leading singular vector orders the conformations along the principal axis of variation in their image likelihoods, and requires neither a trained model nor a reference structure. The image-derived coordinate could replace the user-defined CV in enhanced sampling simulations such that the particle images define the sampling direction.

The prior fails to reach the target distribution at very low SNR, and convergence is therefore driven by the cryo-EM signal rather than by an intrinsic bias in the WE sampling protocol^37,94^. The cryo-EM images are synthetic rather than experimental, and while synthetic images have been shown to be a reasonable proxy for experimental cryo-EM data in prior reweighting studies^77^, the application of *cryoWEight* to experimental datasets remains to be demonstrated. For experimental datasets, the particle pose and CTF are obtained as estimates from consensus refinement, and the likelihood must therefore account for the associated uncertainty. Forward models that marginalize over the pose, the CTF, and pixel masking are available in cryoLike^156^, cryoJAX^157^, and BioEM^74,75^. The dissimilarity metrics are computed relative to a known cryo-EM distribution, which is unavailable for experimental images, and therefore convergence would need to be assessed by a separate criterion.

In summary, *cryoWEight* is an advanced sampling strategy that guides path sampling in WE simulations using cryo-EM images. The framework integrates the complementary strengths of the two approaches, where WE simulations generate new conformations in undersampled regions via splitting and merging operations, and cryo-EM reweighting corrects the population distribution to match experimentally observed ensembles, which then seed further WE sampling until convergence is achieved. The current methodology can be extended to real cryo-EM datasets, in which the conformation weights are determined by the likelihood of each particle image and no separate model of conformational heterogeneity is required. While we have presented benchmarks of this framework on smaller tractable systems for MD simulation, *cryoWEight* offers a framework for physics-based modeling to capture the distribution of conformational states in cryo-EM data, potentially reconstructing large-scale transitions of biomolecular assemblies^158–161^, resolving conformational heterogeneity hidden by current methods that average out distinct conformations.

## Supplementary Material

The supplementary material includes a single *cryoWEight* iteration workflow (Figure S1). Tables S1-S3 provide the parameters for unbiased molecular dynamics simulations, synthetic cryo-EM image generation, and weighted ensemble simulations for the chignolin, NTL9, and ADK proteins. Tables S4, S6, and S7 report the distribution dissimilarity metrics against the cryo-EM target distribution for the chignolin, NTL9, and ADK proteins, respectively, and Tables S5, S9, and S8 report the corresponding effective sample size analyses.

## Acknowledgments

All simulations were performed on the Rusty cluster at the Flatiron Institute, a division of the Simons Foundation. The authors thank Pilar Cossio and Miro Astore for helpful discussions and constructive comments on the manuscript.

## Data Availability Statement

The *cryoWEight* source code is available at https://github.com/anandojha/cryoWEight.

