## Supplementary File for "*cryoWEight*: Conformational ensembles from cryo-EM particle images with weighted ensemble simulations"

### Supplementary information for *cryoWEight*: Conformational ensembles from cryo-EM particle images with weighted ensemble simulations

Anupam Anand Ojha,[1] Praveen Ranganath Prabhakar,[2,3] Ioan Andricioaei,[2,4] and Sonya M. Hanson*[1]

[1]*Center for Computational Biology and Center for Computational Mathematics, Flatiron Institute, New York, NY 10010, USA*
[2]*Department of Chemistry, University of California Irvine, Irvine, CA 92697, USA*
[3]*Current Affiliation: Department of Chemistry, University of Michigan, Ann Arbor, MI 48109, USA*
[4]*Department of Physics and Astronomy, University of California Irvine, Irvine, CA 92697, USA*

*

**Input.** Ensemble $\mathbf{x}aa = 1^T$ of $T$ conformations from the prior MD trajectory ($\ell = 0$) or the merged WE trajectory ($\ell > 0$), where $\ell$ is the iteration index of the protocol, together with $N$ cryo-EM particle images $\mathscr{I}nn = 1^N$, bin width $\Delta$ of the collective variable grid, and convergence tolerance $\varepsilon$. Each particle image carries the contrast transfer function and pixelwise noise of width $\sigma_{\text{noise}} = \sigma_{\text{signal}}/\sqrt{\text{SNR}}$, where $\sigma_{\text{signal}}$ is the root mean square intensity of the noise-free image and SNR is the signal-to-noise ratio.

↓

**Step 1.** Assign the uniform weight $\omega_a^{(\ell)} = 1/T$ to every conformation $a$ and compute its collective variables $\text{CV}a = (q_a^{(1)}, q_a^{(2)})$.

↓

**Step 2.** Bin the conformations on the grid of width $\Delta$, select a representative frame $fu,v$ from the conformations $S_{u,v}$ occupying each bin $(u,v)$, and merge their weights into $\omega' fu,v = \sum_{a\in S_{u,v}} \omega_a^{(\ell)}$ to obtain $M$ representative structures.

↓

**Step 3.** Project each representative onto the image plane with the rotation matrix $\mathbf{R}$ and the contrast transfer function of particle $\mathscr{I}n$, and subtract the mean intensity from the template $I_m(p) = \frac{1}{2\pi\sigma\text{residue}^2 N_{\text{residue}}} \sum_j \exp!\left(-\frac{\|\mathbf{y}p-(\mathbf{R}\mathbf{r}j)xy\|^2}{2\sigma\text{residue}^2}\right)$, where $\sigma_{\text{residue}}$ is the width of the Gaussian placed on each residue, $N_{\text{residue}}$ is the number of residues, $\mathbf{y}p$ is the center of pixel $p$, and $\mathbf{r}j$ is the position of the C$\alpha$ atom of residue $j$.

↓

**Step 4.** Compute the squared distance $dmn^2 = \sum p\,[I_m(p) - \mathscr{I}n(p)]^2$ between the template of structure $m$ and particle $n$, and the log-likelihood $\log Pmn = -d_{mn}^2/2\lambda^2 - (P/2)\log(2\pi\lambda^2)$, where $P$ is the number of pixels and the noise scale $\lambda$ is averaged over one noise-free synthetic image per representative.

↓

**Step 5.** Iterate the expectation-maximization updates

$$r_{mn} \leftarrow \frac{\omega_m P_{mn}}{\sum_{m'} \omega_{m'} P_{m'n}} \quad \text{(E step)} \qquad \omega_m \leftarrow \frac{1}{N}\sum_n r_{mn} \quad \text{(M step)}$$

where $r_{mn}$ is the posterior probability that image $n$ arose from structure $m$, until the weights and the average log-likelihood $\mathscr{L} = \frac{1}{N}\sum_n \log \sum_m \omega_m P_{mn}$ stabilize at the optimized weights $\omega_m^*$.

↓

**Step 6.** Redistribute the optimized weight of each representative to the conformations it represents, $\omega_a^* = \omega_{fu,v}^*/|S_{u,v}|$ for $a \in S_{u,v}$, and write the reweighted ensemble $\{\mathbf{x}_m, \omega_m^*\}$ as the starting structures for the next WE simulation.

↓

**Step 7.** Compute the frame density $\rho u,v$, the fraction of prior conformations in bin $(u,v)$, and place the adaptive binning anchor at the occupied bin of highest density at least $k$ standard deviations from the prior mean along the progress coordinate $\xi$, on the side facing the target, relaxing $k = 3,2,1$ until occupied bins remain.

↓

**Step 8.** Run $N_{\text{WE}}$ WE subiterations, each of length $\tau$, splitting and merging walkers on the discretized CV space.

↓

**Step 9.** Combine the segment trajectories into the ensemble of iteration $\ell+1$ and project it onto the CV space to obtain the binned probability distribution $P^{(\ell+1)} = P_{u,v}$.

↓

**Step 10.** Stop once the Kullback-Leibler divergence falls below the tolerance, $D_{\text{KL}}!\left(P^{(\ell+1)}|P^{(\ell)}\right) < \varepsilon$, otherwise return to Step 1 with the ensemble of iteration $\ell+1$.

FIG. S1: Workflow of a single *cryoWEight* iteration. The input ensemble is clustered into representative structures, the representatives are reweighted against the cryo-EM particle images, and the reweighted conformations seed the next weighted ensemble simulation. The cycle repeats until the Kullback-Leibler divergence between consecutive iterations falls below the user-defined tolerance.

| a) MD simulation parameters | | | |
|---|---|---|---|
| Force field | Amber ff14SB | Water model | TIP3P |
| Ensemble | NVT | Temperature | 300 K |
| Langevin friction | 1 $ps^{-1}$ | Integration timestep | 2 fs |
| Particle-mesh Ewald cutoff | 1.0 nm | Hydrogen bond constraint | On |
| Protein residues | 10 | Protein atoms | 138 |
| Water molecules | 799 | Water atoms | 2397 |
| Total system size | 2535 atoms | Cubic box edge | 31.33 Å |
| Counterions | None added | | |
| **b) Image generation parameters** | | | |
| Image size | 128×128 | Pixel size | 0.2 Å |
| Residue Gaussian width ($\sigma_{\text{residue}}$) | 1.5 Å | Images per structure | 1 |
| SNR mask radius | $0.4\,n_{\text{pixel}}$ | | |
| Contrast transfer function | Enabled | Defocus | [0.027 µm, 0.090 µm] |
| Expectation-maximization iterations | 5000 | Target number of images | 10000 |
| **c) WE simulation parameters** | | | |
| Resampling time per WE subiteration ($\tau$) | 20 ps | Walkers per bin | 4 |
| WE subiterations per *cryoWEight* iteration ($N_{\text{WE}}$) | 30 | Minimal adaptive binning grid | 15×15 |

d) Convergence statistics

| i) MAB anchor placement at SNR = 1.0 | | | | ii) SNR dependence at distal MAB anchor | | | |
|---|---|---|---|---|---|---|---|
| MAB anchor ($\|k\|$) | Iterations | Sampling (µs) | Converged | SNR | Iterations | Sampling (µs) | Converged |
| Distal ($\|k\| = 3$) | 4 | 1.48 | Yes | 1.0 | 4 | 1.48 | Yes |
| Medial ($\|k\| = 2$) | 4 | 1.82 | Yes | $10^{-2}$ | 6 | 2.79 | Yes |
| Proximal ($\|k\| = 1$) | 8 | 3.38 | Yes | $10^{-8}$ | 8 | 3.38 | No |

TABLE S1: Simulation parameters and convergence statistics for *cryoWEight* simulations on chignolin protein. Panels a, b, and c report parameters for MD simulations, image generation, and WE simulations, respectively. Panel d reports the iterations and sampling time required for *cryoWEight* simulations as a function of (i) the MAB anchor placement at SNR = 1.0, and (ii) the SNR at the distal MAB anchor placement.

| a) MD simulation parameters | | | |
|---|---|---|---|
| Force field | Amber ff14SB | Solvent model | OBC2 implicit |
| Ensemble | NVT | Temperature | 300 K |
| Langevin friction | 5 $ps^{-1}$ | Integration timestep | 2 fs |
| Non-bonded cutoff | 2.0 nm | Hydrogen bond constraint | On |
| Solvent dielectric | 78.5 | Solute dielectric | 1.0 |
| Protein residues | 39 | Protein atoms | 627 |
| Total system size | 627 atoms | Periodic boundaries | None |
| **b) Image generation parameters** | | | |
| Image size | 128×128 | Pixel size | 0.5 Å |
| Residue Gaussian width ($\sigma_{\text{residue}}$) | 1.5 Å | Images per structure | 1 |
| SNR mask radius | $0.4\,n_{\text{pixel}}$ | | |
| Contrast transfer function | Enabled | Defocus | [0.027 µm, 0.090 µm] |
| Expectation-maximization iterations | 5000 | Target number of images | 10000 |
| **c) WE simulation parameters** | | | |
| Resampling time per WE subiteration ($\tau$) | 40 ps | Walkers per bin | 4 |
| WE subiterations per *cryoWEight* iteration ($N_{\text{WE}}$) | 40 | Minimal adaptive binning grid | 18×18 |

d) Convergence statistics

| i) SNR dependence (target RMSD $\in [2.0, 3.0]$ Å) | | | | ii) Target dependence (SNR = 1.0) | | | |
|---|---|---|---|---|---|---|---|
| SNR | Iterations | Sampling (µs) | Converged | RMSD (Å) | Iterations | Sampling (µs) | Converged |
| 1.0 | 4 | 6.45 | Yes | $[2.00, 3.00]$ | 4 | 6.45 | Yes |
| $10^{-8}$ | 8 | 15.76 | No | $[2.75, 3.50]$ | 3 | 4.58 | Yes |

TABLE S2: Simulation parameters and convergence statistics for *cryoWEight* simulations on NTL9 protein. Panels a, b, and c report parameters for MD simulations, image generation, and WE simulations, respectively. Panel d reports the iterations and sampling time required for *cryoWEight* simulations as a function of (i) the SNR at the distal MAB anchor with target RMSD $\in [2.0, 3.0]$ Å, and (ii) the target state at SNR = 1.0 with the distal MAB anchor.

| a) MD simulation parameters | | | |
|---|---|---|---|
| Force field | Amber ff14SB | Water model | TIP3P |
| Ensemble | NVT | Temperature | 300 K |
| Langevin friction | 1 $ps^{-1}$ | Integration timestep | 2 fs |
| Particle-mesh Ewald cutoff | 1.0 nm | Hydrogen bond constraint | On |
| Protein residues | 214 | Protein atoms | 3341 |
| Water molecules | 9264 | Water atoms | 27 792 |
| Total system size | 31 137 atoms | Cubic box edge | 69.16 Å |
| Counterions | 4 $Na^{+}$ | | |
| **b) Image generation parameters** | | | |
| Image size | 200×200 | Pixel size | 0.5 Å |
| Residue Gaussian width ($\sigma_{\text{residue}}$) | 1.5 Å | Images per structure | 1 |
| SNR mask radius | $0.4\,n_{\text{pixel}}$ | | |
| Contrast transfer function | Enabled | Defocus | [0.027 μm, 0.090 μm] |
| Expectation-maximization iterations | 5000 | Target number of images | 2500 |
| **c) WE simulation parameters** | | | |
| Resampling time per WE subiteration ($\tau$) | 100 ps | Walkers per bin | 4 |
| WE subiterations per *cryoWEight* iteration ($N_{\text{WE}}$) | 25 | Minimal adaptive binning grid | 15×15 |

TABLE S3: Simulation parameters for *cryoWEight* simulations on ADK protein. Panels a, b, and c report parameters for MD simulations, image generation, and WE simulations, respectively.

| Iteration | Kullback-Leibler divergence | Jensen-Shannon divergence | Wasserstein distance (Å) | Bhattacharyya distance |
|---|---|---|---|---|
| SNR = 1.0 at distal MAB anchor | | | | |
| Initial ensemble | 21.513 | 0.693 | 6.433 | 10.133 |
| Iteration 1 | 21.508 | 0.693 | 3.107 | 9.719 |
| Iteration 2 | 16.961 | 0.691 | 1.658 | 5.061 |
| Iteration 3 | 3.191 | 0.445 | 0.264 | 0.823 |
| Iteration 4 | 0.357 | 0.108 | 0.083 | 0.144 |
| SNR = $10^{-2}$ at distal MAB anchor | | | | |
| Initial ensemble | 21.513 | 0.693 | 6.393 | 10.133 |
| Iteration 1 | 14.784 | 0.693 | 3.052 | 5.388 |
| Iteration 2 | 5.340 | 0.674 | 2.109 | 2.587 |
| Iteration 3 | 4.201 | 0.647 | 1.651 | 2.034 |
| Iteration 4 | 4.146 | 0.642 | 1.289 | 1.977 |
| Iteration 5 | 3.708 | 0.625 | 1.028 | 1.781 |
| Iteration 6 | 0.308 | 0.092 | 0.085 | 0.122 |
| SNR = $10^{-8}$ at distal MAB anchor | | | | |
| Initial ensemble | 21.513 | 0.693 | 6.358 | 10.133 |
| Iteration 1 | 10.732 | 0.690 | 2.940 | 4.112 |
| Iteration 2 | 21.513 | 0.693 | 4.386 | 9.927 |
| Iteration 3 | 21.513 | 0.693 | 4.302 | 9.822 |
| Iteration 4 | 14.202 | 0.691 | 3.339 | 4.885 |
| Iteration 5 | 15.966 | 0.691 | 3.448 | 5.136 |
| Iteration 6 | 14.417 | 0.691 | 3.437 | 4.763 |
| Iteration 7 | 10.971 | 0.690 | 3.228 | 4.169 |
| Iteration 8 | 9.273 | 0.686 | 2.968 | 3.584 |
| SNR = 1.0 at medial MAB anchor | | | | |
| Initial ensemble | 21.513 | 0.693 | 6.391 | 10.133 |
| Iteration 1 | 20.254 | 0.693 | 3.480 | 6.986 |
| Iteration 2 | 9.420 | 0.686 | 2.071 | 3.529 |
| Iteration 3 | 4.425 | 0.650 | 1.034 | 2.093 |
| Iteration 4 | 0.310 | 0.092 | 0.067 | 0.120 |
| SNR = 1.0 at proximal MAB anchor | | | | |
| Initial ensemble | 21.513 | 0.693 | 6.548 | 10.133 |
| Iteration 1 | 21.499 | 0.693 | 3.834 | 9.564 |
| Iteration 2 | 21.034 | 0.693 | 2.877 | 8.781 |
| Iteration 3 | 8.426 | 0.679 | 1.962 | 3.060 |
| Iteration 4 | 7.443 | 0.675 | 1.421 | 2.849 |
| Iteration 5 | 7.312 | 0.664 | 1.170 | 2.558 |
| Iteration 6 | 5.334 | 0.584 | 0.461 | 1.551 |
| Iteration 7 | 2.839 | 0.438 | 0.158 | 0.768 |
| Iteration 8 | 0.322 | 0.092 | 0.058 | 0.116 |

TABLE S4: Distribution dissimilarity metrics between the sampled conformational ensemble and the cryo-EM distribution for the chignolin protein across successive iterations for (i) SNR dependence at the distal MAB anchor ($|k| = 3$) at SNR = 1.0, $10^{-2}$, and $10^{-8}$, and (ii) MAB anchor placement dependence at SNR = 1.0 for distal ($|k| = 3$), medial ($|k| = 2$), and proximal ($|k| = 1$) anchor locations.

| Iteration | Simulation time ($\mu$s) | Frames (N) | Effective sample size (ESS) | ESS/N (%) |
|---|---|---|---|---|
| SNR = 1.0 at distal MAB anchor | | | | |
| Initial ensemble | 0.20 | 210 | 72 | 34.13 |
| Iteration 1 | 0.34 | 548 | 18 | 3.28 |
| Iteration 2 | 0.42 | 846 | 65 | 7.72 |
| Iteration 3 | 0.12 | 927 | 101 | 10.88 |
| Iteration 4 | 0.40 | 1305 | 114 | 8.73 |
| SNR = $10^{-2}$ at distal MAB anchor | | | | |
| Initial ensemble | 0.20 | 210 | 54 | 25.92 |
| Iteration 1 | 0.43 | 687 | 21 | 3.11 |
| Iteration 2 | 0.50 | 1142 | 78 | 6.82 |
| Iteration 3 | 0.48 | 1199 | 104 | 8.70 |
| Iteration 4 | 0.44 | 1276 | 118 | 9.27 |
| Iteration 5 | 0.31 | 1202 | 125 | 10.42 |
| Iteration 6 | 0.43 | 1256 | 134 | 10.71 |
| SNR = $10^{-8}$ at distal MAB anchor | | | | |
| Initial ensemble | 0.20 | 210 | 210 | 100.00 |
| Iteration 1 | 0.35 | 620 | 620 | 100.00 |
| Iteration 2 | 0.37 | 470 | 460 | 97.96 |
| Iteration 3 | 0.38 | 547 | 547 | 100.00 |
| Iteration 4 | 0.41 | 830 | 724 | 87.23 |
| Iteration 5 | 0.42 | 879 | 795 | 90.47 |
| Iteration 6 | 0.41 | 1055 | 961 | 91.04 |
| Iteration 7 | 0.42 | 1136 | 987 | 86.90 |
| Iteration 8 | 0.42 | 1129 | 1129 | 100.00 |
| SNR = 1.0 at medial MAB anchor | | | | |
| Initial ensemble | 0.20 | 210 | 69 | 32.82 |
| Iteration 1 | 0.42 | 574 | 26 | 4.46 |
| Iteration 2 | 0.29 | 897 | 59 | 6.59 |
| Iteration 3 | 0.48 | 1205 | 96 | 7.93 |
| Iteration 4 | 0.43 | 1214 | 141 | 11.59 |
| SNR = 1.0 at proximal MAB anchor | | | | |
| Initial ensemble | 0.20 | 210 | 74 | 35.43 |
| Iteration 1 | 0.42 | 568 | 20 | 3.46 |
| Iteration 2 | 0.47 | 712 | 57 | 8.03 |
| Iteration 3 | 0.46 | 1088 | 76 | 7.02 |
| Iteration 4 | 0.44 | 1183 | 90 | 7.65 |
| Iteration 5 | 0.18 | 1088 | 89 | 8.18 |
| Iteration 6 | 0.50 | 1174 | 70 | 5.97 |
| Iteration 7 | 0.25 | 1291 | 132 | 10.20 |
| Iteration 8 | 0.46 | 1302 | 138 | 10.63 |

TABLE S5: Effective sample size analysis for the chignolin protein across successive iterations for (i) SNR dependence at the distal MAB anchor ($|k| = 3$) at SNR = 1.0, $10^{-2}$, and $10^{-8}$, and (ii) MAB anchor placement dependence at SNR = 1.0 for distal ($|k| = 3$), medial ($|k| = 2$), and proximal ($|k| = 1$) anchor locations.

| Iteration | Kullback-Leibler divergence | Jensen-Shannon divergence | Wasserstein distance (Å) | Bhattacharyya distance |
|---|---|---|---|---|
| SNR = 1.0 at distal MAB anchor and target RMSD $\in$ [2.0, 3.0] Å | | | | |
| Initial ensemble | 22.350 | 0.693 | 5.103 | 10.103 |
| Iteration 1 | 22.350 | 0.693 | 2.418 | 10.129 |
| Iteration 2 | 18.831 | 0.692 | 1.386 | 5.930 |
| Iteration 3 | 6.503 | 0.605 | 0.466 | 1.759 |
| Iteration 4 | 1.338 | 0.295 | 0.097 | 0.426 |
| SNR = $10^{-8}$ at distal MAB anchor and target RMSD $\in$ [2.0, 3.0] Å | | | | |
| Initial ensemble | 22.918 | 0.693 | 4.435 | 10.185 |
| Iteration 1 | 22.918 | 0.693 | 3.969 | 10.021 |
| Iteration 2 | 22.918 | 0.693 | 3.971 | 9.938 |
| Iteration 3 | 22.918 | 0.693 | 3.944 | 9.925 |
| Iteration 4 | 22.585 | 0.693 | 3.687 | 9.708 |
| Iteration 5 | 22.651 | 0.693 | 3.837 | 9.712 |
| Iteration 6 | 22.765 | 0.693 | 3.713 | 9.740 |
| Iteration 7 | 22.178 | 0.693 | 3.819 | 9.603 |
| Iteration 8 | 19.065 | 0.693 | 3.808 | 8.589 |
| SNR = 1.0 at distal MAB anchor and target RMSD $\in$ [2.75, 3.50] Å | | | | |
| Initial ensemble | 22.918 | 0.693 | 4.426 | 10.185 |
| Iteration 1 | 22.918 | 0.693 | 1.755 | 10.233 |
| Iteration 2 | 9.154 | 0.688 | 0.821 | 3.813 |
| Iteration 3 | 1.474 | 0.365 | 0.179 | 0.621 |

TABLE S6: Distribution dissimilarity metrics between the sampled conformational ensemble and cryo-EM distribution for the NTL9 protein across successive iterations for (i) SNR dependence at the distal MAB anchor ($|k| = 3$) at SNR = 1.0 and $10^{-8}$ for target RMSD $\in$ [2.0, 3.0] Å, and (ii) target state dependence at the distal MAB anchor ($|k| = 3$) at SNR = 1.0 for target RMSD $\in$ [2.0, 3.0] Å and $\in$ [2.75, 3.50] Å.

| Iteration | Kullback-Leibler divergence | Jensen-Shannon divergence | Wasserstein distance (°) | Bhattacharyya distance |
|---|---|---|---|---|
| Initial ensemble | 20.710 | 0.693 | 20.429 | 8.661 |
| Iteration 1 | 19.475 | 0.693 | 19.938 | 6.704 |
| Iteration 2 | 3.445 | 0.603 | 10.256 | 1.597 |
| Iteration 3 | 2.469 | 0.514 | 7.243 | 1.094 |
| Iteration 4 | 2.268 | 0.487 | 6.627 | 0.983 |
| Iteration 5 | 2.260 | 0.483 | 6.666 | 0.967 |
| Iteration 6 | 2.266 | 0.483 | 6.817 | 0.970 |
| Iteration 7 | 2.280 | 0.484 | 6.812 | 0.974 |

TABLE S7: Distribution dissimilarity metrics between the sampled conformational ensemble and cryo-EM distribution ($\theta_{\text{NMP}} \in [70°, 85°]$, $\theta_{\text{LID}} \in [130°, 150°]$) for the ADK protein across successive iterations at SNR = 1.0 with the proximal MAB anchor ($|k| = 1$).

| Iteration | Simulation time ($\mu$s) | Frames (N) | Effective sample size (ESS) | ESS/N (%) |
|---|---|---|---|---|
| Initial ensemble | 0.02 | 117 | 52 | 44.23 |
| Iteration 1 | 1.89 | 686 | 16 | 2.28 |
| Iteration 2 | 2.17 | 1530 | 78 | 5.11 |
| Iteration 3 | 1.72 | 1783 | 49 | 2.73 |
| Iteration 4 | 1.74 | 1867 | 57 | 3.05 |
| Iteration 5 | 1.89 | 1945 | 91 | 4.69 |
| Iteration 6 | 1.91 | 1918 | 96 | 5.02 |
| Iteration 7 | 1.92 | 2003 | 127 | 6.34 |

TABLE S8: Effective sample size analysis for the ADK protein across successive iterations at SNR = 1.0 with the proximal MAB anchor ($|k| = 1$), with cryo-EM target distribution defined by $\theta_{\text{NMP}} \in [70°, 85°]$ and $\theta_{\text{LID}} \in [130°, 150°]$.

| Iteration | Simulation time ($\mu$s) | Frames (N) | Effective sample size (ESS) | ESS/N (%) |
|---|---|---|---|---|
| SNR = 1.0 at distal MAB anchor and target RMSD ∈ [2.0, 3.0] Å | | | | |
| Initial ensemble | 0.20 | 364 | 108 | 29.57 |
| Iteration 1 | 1.39 | 1669 | 193 | 11.59 |
| Iteration 2 | 1.71 | 2271 | 135 | 5.93 |
| Iteration 3 | 1.75 | 2513 | 199 | 7.91 |
| Iteration 4 | 1.40 | 3241 | 61 | 1.89 |
| SNR = $10^{-8}$ at distal MAB anchor and target RMSD ∈ [2.0, 3.0] Å | | | | |
| Initial ensemble | 0.20 | 364 | 364 | 100.00 |
| Iteration 1 | 1.67 | 1699 | 1697 | 99.86 |
| Iteration 2 | 1.90 | 2191 | 2188 | 99.88 |
| Iteration 3 | 1.96 | 2317 | 2292 | 98.92 |
| Iteration 4 | 1.98 | 2636 | 2635 | 99.96 |
| Iteration 5 | 2.02 | 2929 | 2880 | 98.34 |
| Iteration 6 | 1.99 | 2917 | 2862 | 98.11 |
| Iteration 7 | 2.01 | 2977 | 2917 | 97.98 |
| Iteration 8 | 2.03 | 3025 | 2967 | 98.08 |
| SNR = 1.0 at distal MAB anchor and target RMSD ∈ [2.75, 3.50] Å | | | | |
| Initial ensemble | 0.20 | 364 | 100 | 27.47 |
| Iteration 1 | 1.46 | 1720 | 126 | 7.32 |
| Iteration 2 | 1.54 | 2519 | 153 | 6.09 |
| Iteration 3 | 1.38 | 2456 | 165 | 6.73 |

TABLE S9: Effective sample size analysis for the NTL9 protein across successive iterations for (i) SNR dependence at the distal MAB anchor ($|k| = 3$) at SNR = 1.0 and $10^{-8}$ for target RMSD ∈ [2.0, 3.0] Å, and (ii) target state dependence at the distal MAB anchor ($|k| = 3$) at SNR = 1.0 for target RMSD ∈ [2.0, 3.0] Å and ∈ [2.75, 3.50] Å.

#### Glossary

| Term | Definition / meaning | Term | Definition |
|---|---|---|---|
| **Acronyms** | | | |
| ADK | Adenylate kinase | NMP | Nucleoside monophosphate |
| ADP | Adenosine diphosphate | NMR | Nuclear magnetic resonance |
| AMP | Adenosine monophosphate | NTL9 | N-terminal domain of ribosomal protein L9 |
| ATP | Adenosine triphosphate | NVT | Canonical ensemble (constant $N$, $V$, $T$) |
| CHARMM | Chemistry at Harvard Macromolecular Mechanics | OBC2 | Onufriev-Bashford-Case generalized Born model 2 |
| CORE | Rigid central domain of adenylate kinase | OpenMM | Open Molecular Mechanics simulation toolkit |
| cryo-EM | Cryogenic electron microscopy | PDB | Protein Data Bank |
| CTF | Contrast transfer function | PME | Particle mesh Ewald |
| CV | Collective variable | RCSB | Research Collaboratory for Structural Bioinformatics |
| MD | Molecular dynamics | RMSD | Root mean square deviation |
| ESS | Effective sample size | $R_g$ | Radius of gyration |
| FE | Free energy | SAXS | Small-angle X-ray scattering |
| JS | Jensen-Shannon divergence | SNR | Signal-to-noise ratio |
| KL | Kullback-Leibler divergence | TIP3P | Three-site transferable intermolecular potential |
| LID | ATP-binding lid domain of adenylate kinase | WE | Weighted ensemble |
| MAB | Minimal adaptive binning | WESTPA | Weighted Ensemble Simulation Toolkit with Parallelization and Analysis |
| **Symbols: Weighted ensemble simulations** | | | |
| $\Omega$ | Configuration space | $B_j$ | The $j$-th bin in CV space |
| $M_\mathrm{w}$ | Total number of walkers (WE) | $n_1, n_2$ | Number of bins along each CV dimension |
| $K$ | Number of distinct initial configurations | $j_1, j_2$ | Bin indices along each CV dimension |
| $\mathbf{X}_k$ | The $k$-th initial configuration | $a_{j_1}, a'_{j_1}$ | Lower / upper bin boundaries in the first coordinate |
| $p_k$ | Initial probability assigned to configuration $k$ | $b_{j_2}, b'_{j_2}$ | Lower / upper bin boundaries in the second coordinate |
| $M_k$ | Walkers spawned from configuration $k$ | $S_j^{(n)}$ | Set of walkers in bin $B_j$ at iteration $n$ |
| $w_i^{(n)}$ | Statistical weight of walker $i$ at WE iteration $n$ | $W_j^{(n)}$ | Total bin weight, $\sum_{i\in S_j^{(n)}} w_i^{(n)}$ |
| $P^{(0)}(\mathbf{X})$ | Initial probability distribution over configurations | $\tau$ | WE resampling interval |
| $\delta(\cdot)$ | Dirac delta function | $\mathscr{D}_\tau$ | Stochastic propagator over interval $\tau$ |
| $\mathbf{q}$ | Collective variable vector ($d$-dimensional) | $M_\mathrm{bin}$ | Target number of walkers per bin |
| $d$ | Dimensionality of CV space | $m_i$ | Number of split copies of walker $i$ |
| $N$ | Number of bins in CV space | $k_B\mathscr{T}$ | Thermal energy ($k_B$ Boltzmann constant, $\mathscr{T}$ temperature) |
| **Symbols: cryo-EM ensemble reweighting** | | | |
| $T$ | Number of MD frames in the input trajectory | $P$ | Number of pixels per image |
| $\mathbf{x}_a$ | The $a$-th MD frame | $d^2_{mn}$ | Pixelwise squared distance between $I_m$, $\mathscr{I}_n$ |
| $\omega_a^{(0)}$ | Uniform initial frame weight, $1/T$ | $\lambda$ | Noise standard deviation of the Gaussian likelihood |
| $\mathrm{CV}_a$ | Collective variables of frame $a$ | $\log P_{mn}$ | Gaussian log-likelihood for pair $(m,n)$ |
| $\Delta$ | Bin width in CV space (clustering grid) | $t$ | Expectation-maximization iteration index |
| $N_1, N_2$ | Number of clustering bins along each dimension | $r_{mn}^{(t)}$ | Posterior probability that image $n$ was generated by structure $m$ |
| $(u,v)$ | Bin index pair in the 2D CV grid | $m'$ | Dummy structure index in the expectation-maximization normalization |
| $S_{u,v}$ | Set of frame indices in bin $(u,v)$ | $\omega_m^{(t)}$ | Structure weight at expectation-maximization iteration $t$ |
| $f_{u,v}$ | Representative frame for bin $(u,v)$ | $\mathscr{L}^{(t)}$ | Average log-likelihood at expectation-maximization iteration $t$ |
| $\omega'_a$, $\omega''_a$ | Bin-merged and renormalized frame weights | $\omega_m^*$ | Optimized structure weight at expectation-maximization convergence |
| $M$ | Number of structures in occupied bins | $\sigma_\mathrm{residue}$ | Width of the Gaussian placed on each residue |
| $\mathbf{x}_m$ | The $m$-th representative structure | $\mathbf{r}_j$ | Position of the C$\alpha$ atom of residue $j$ |
| $N$ | Number of cryo-EM images | $N_\mathrm{residue}$ | Number of residues |
| $\mathscr{I}_n$ | The $n$-th cryo-EM image | $\mathbf{R}$ | Rotation matrix defining the projection direction |
| $I_m(p)$ | Pixel $p$ of the template from $\mathbf{x}_m$ | $\mathbf{y}_p$ | Center of pixel $p$ |
| $\sigma_\mathrm{signal}$ | Root mean square intensity of the noise-free image | $\sigma_{\mathrm{noise},m}$ | Noise standard deviation for representative structure $m$ |
| **Symbols: Iterative *cryoWEight* protocol and convergence** | | | |
| $P_{u,v}$ | Weighted probability of bin $(u,v)$ | $P^{(\ell)}$ | Binned distribution at iteration $\ell$ |
| $G_{u,v}$ | Free energy of bin $(u,v)$ | $N_\mathrm{WE}$ | WE subiterations per *cryoWEight* iteration |
| $(u^*, v^*)$ | MAB anchor bin indices | $\theta_\mathrm{NMP}$, $\theta_\mathrm{LID}$ | ADK NMP- and LID-domain opening angles |
| $\ell$ | *cryoWEight* iteration index (outer loop) | ESS | Effective sample size, $1/\sum_i w_i^2$ |
| $\varepsilon$ | Tolerance on inter-iteration KL divergence | $\{w_i\}$ | Normalized frame weights |
| $D_\mathrm{KL}(\cdot\|\cdot)$ | Kullback-Leibler divergence | $k$ | Anchor cutoff in standard deviations of the prior along the progress coordinate |
